# EXPERT: Transfer Learning-enabled context-aware microbial source tracking

**DOI:** 10.1101/2021.01.29.428751

**Authors:** Hui Chong, Qingyang Yu, Yuguo Zha, Guangzhou Xiong, Nan Wang, Xinhe Huang, Shijuan Huang, Chuqing Sun, Sicheng Wu, Wei-Hua Chen, Luis Pedro Coelho, Kang Ning

## Abstract

Microbial source tracking quantifies the potential origin of microbial communities, facilitates better understanding of how the taxonomic structure and community functions were formed and maintained. However, previous methods involve a tradeoff between speed and accuracy, and have encountered difficulty in source tracking under many context-dependent settings. Here, we present EXPERT for context-aware microbial source tracking, in which we adopted a Transfer Learning approach to profoundly elevate and expand the applicability of source tracking, enabling biologically informed novel microbial knowledge discovery. We demonstrate that EXPERT can predict microbial sources with performance superior to other methods in efficiency and accuracy. More importantly, we demonstrate EXPERT’s context-aware ability on several applications, including tracking the progression of infant gut microbiome development and monitoring the changes of gut microbiome for colorectal cancer patients. Broadly, transfer learning enables accurate and context-aware microbial source tracking and has the potential for novel microbial knowledge discovery.

## Introduction

Millions of microbial communities have been accumulated from hundreds of environments (also known as “biomes”) worldwide^1–3^, which continuously complete the grand picture of the microbiome world, revolutionizing our understanding of the roles microbes play in human health and disease^4^, biogeochemical cycling^5^ and other processes. The relationships between microbial community samples and their biomes, on the other hand, are extremely complicated, owing to the highly dynamic nature of microbial communities and our limited understanding of how they function. Microbial source tracking (MST) quantifies the potential origin of microbial communities, thereby could help us to understand how the taxonomic structure and community functions were formed and maintained^6^. In previous studies, MST has been widely used to quantify the contamination present in (1) environment^7^ and (2) host driven by the contact of host and environment^8^ (e.g., human skin and exposed environments).

MST has the potential to go beyond the scope of microbial contamination in a variety of contexts, including estimating microbial restoration of cesarean-born infants^9^, quantifying the microbial community differences across diseases, as well as characterizing the gut microbial communities of patients during cancer progression. In these contexts, MST could reveal dynamic patterns of microbial communities, and provide insights into the effect of microbial communities on health care for newborns, chronic diseases, and cancer.

Current methods for source tracking, though having made substantial contributions, have limitations in accuracy and scope of application. SourceTracker^10^ and FEAST^9^ model the query (sink) community as a mixture of sources, and estimate source contribution through Markov Chain Monte Carlo (MCMC) and Expectation-Maximization (EM), which, however, leads to a tradeoff between running time and accuracy, and source tracking among thousands of sources take hours^9,10^. Also, these two methods are heavily dependent on reference samples, leading to the needs of manually selecting possible source samples, rendering source tracking incomplete and error-prone. On the other hand, Random Forest^11^ and ONN4MST^10^ utilize supervised learning models for source contribution estimation^6^ but are constrained by adaptivity: Models cannot be directly applied to other MST tasks once they’ve been built for specific context.

To address these limitations, we developed EXPERT, a method based on an adaptive Neural Network (NN) framework and Transfer Learning, for solving the MST problem. Previous studies have shown that Transfer Learning can significantly expand the applicability of supervised learning models^12^. Here Transfer Learning is used to introduce existing knowledge learnt from other microbial samples to diverse contexts and to facilitate source tracking in a context-aware manner. Systematic assessments have shown EXPERT’s capability of quantifying the potential contribution of sources in a fast and accurate manner, and adapting source tracking in different context-dependent settings. More importantly, we demonstrate the utility of EXPERT in several representative contexts, including tracking the development of the infant gut microbiome, as well as tracking the progression of gut microbiome changes in patients with colorectal cancer (CRC). Transfer Learning-enabled EXPERT, we reasoned, could make it easier to discover novel microbial knowledge across a wide range of applications in these contexts.

## Results

### Rationale, adaptive modeling, and multi-faceted applications of EXPERT

EXPERT is a context-aware method for MST that employs both the adaptive NN and Transfer Learning^12^ frameworks, enabling knowledge transfer of MST models. The adaptive NN framework constructs MST models according to a given MST task (**Methods, Supplementary Fig. S1**). Together with Transfer Learning, EXPERT can automatically construct MST models and utilize the knowledge of fundamental models (i.e., existing MST models) to aid in the learning of the newly constructed models. In our study, three fundamental models were introduced for knowledge transfer (**Supplementary Table S1-S5**): the general model (GM, trained and validated on 118,592 communities from 131 representative biomes), the human model (HM, trained and validated on 52,537 communities from 27 human-associated biomes), and the disease model (DM, trained and validated on 13,642 fecal communities from patients of 19 diseases and healthy controls). Additionally, EXPERT utilizes Multi-task Learning^13^, which enables hierarchical MST (**Methods, Supplementary Fig. S1-S2**).

The knowledge transfer process of EXPERT is illustrated in **Fig. 1a**. EXPERT adopted the rationale of Transfer Learning^12^, allowing context-aware MST through three steps, namely transfer, adaptation, and fine-tuning: In the transfer step, EXPERT adapts the fundamental model to an MST context; in the adaptation and fine-tuning steps, EXPERT optimizes the parameters (**Methods, Supplementary Note 1**). The contextualized model can serve a broad-spectrum of source tracking applications (**Fig. 1b**)

**Figure 1:**
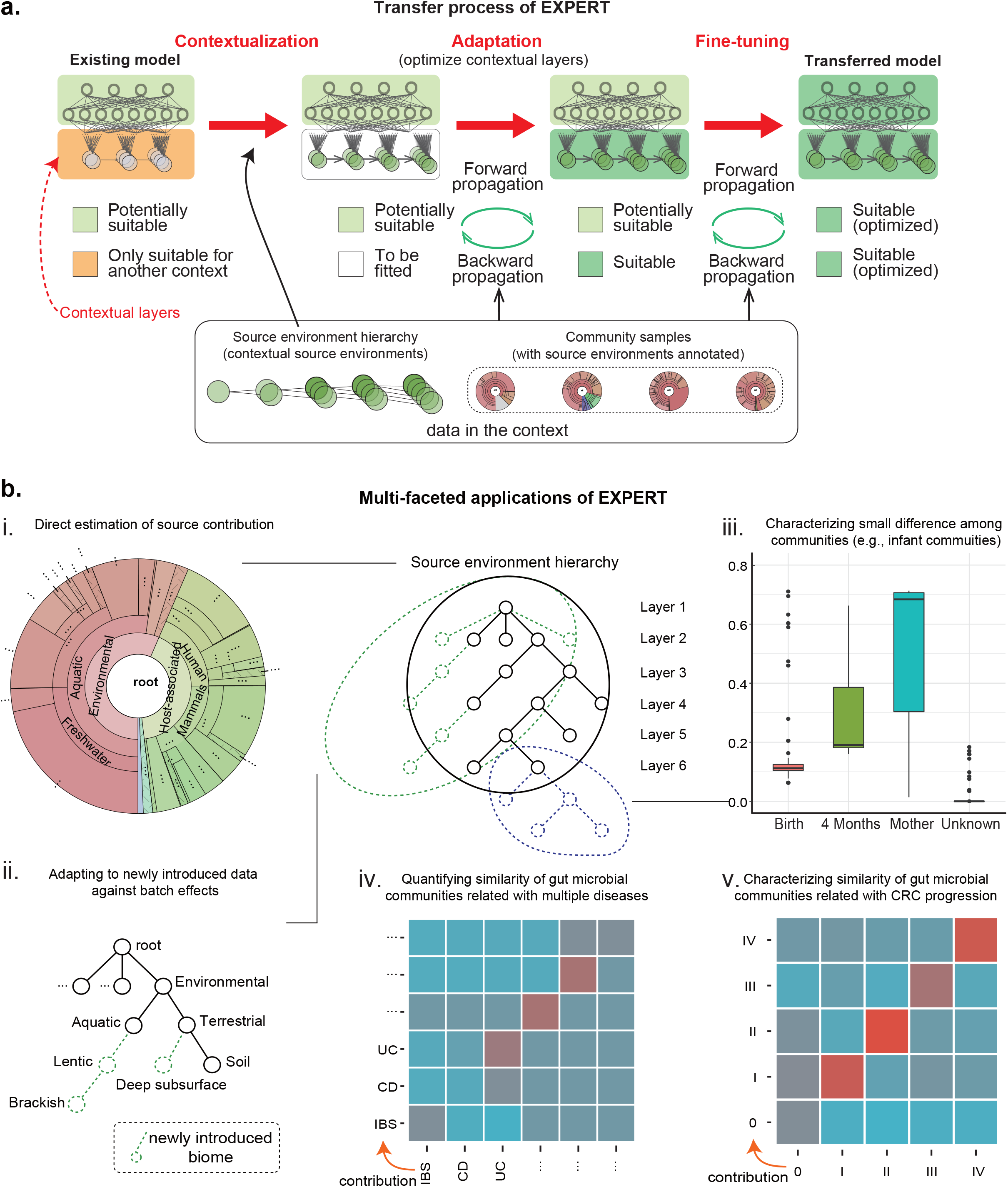
Illustration of EXPERT’s knowledge transfer process. **a**. EXPERT can adapt the knowledge of a fundamental model to an MST context through three steps: transfer (reuse parameters of a fundamental model and reinitialize contextual layers according to the context, red dotted arrows), adaptation (quickly optimize only the contextual layers using iterative forward-backward propagation, green circular arrows), and fine-tuning (further optimize the entire model using the iterative forward-backward propagation). The fundamental model is a pre-trained EXPERT model to be adapted, with several NN layers relatively independent to contexts and a series of contextual NN layers highly specified to a context). Different background colors of the model indicate the suitability of different modules to the context. The contextualized model can serve a broad-spectrum of source tracking applications (based on research purposes, illustrated in **Fig. 1b**). Abbreviations: MST: microbial source tracking; NN: Neural Network.

### Efficiency, accuracy, and adaptivity of EXPERT

Benchmark tests have demonstrated EXPERT’s superior efficiency, accuracy, and adaptivity for MST (**Fig. 2**). Specifically, it outperforms Sourcetracker^6^ and FEAST^9^ in terms of efficiency, while outperforming the NN approach^10^ in terms of accuracy and adaptability. In this part, we assessed these capabilities using 52,537 communities from 27 human-associated biomes (**Supplementary Table S1, S3, Fig. S3**).

**Figure 2.**
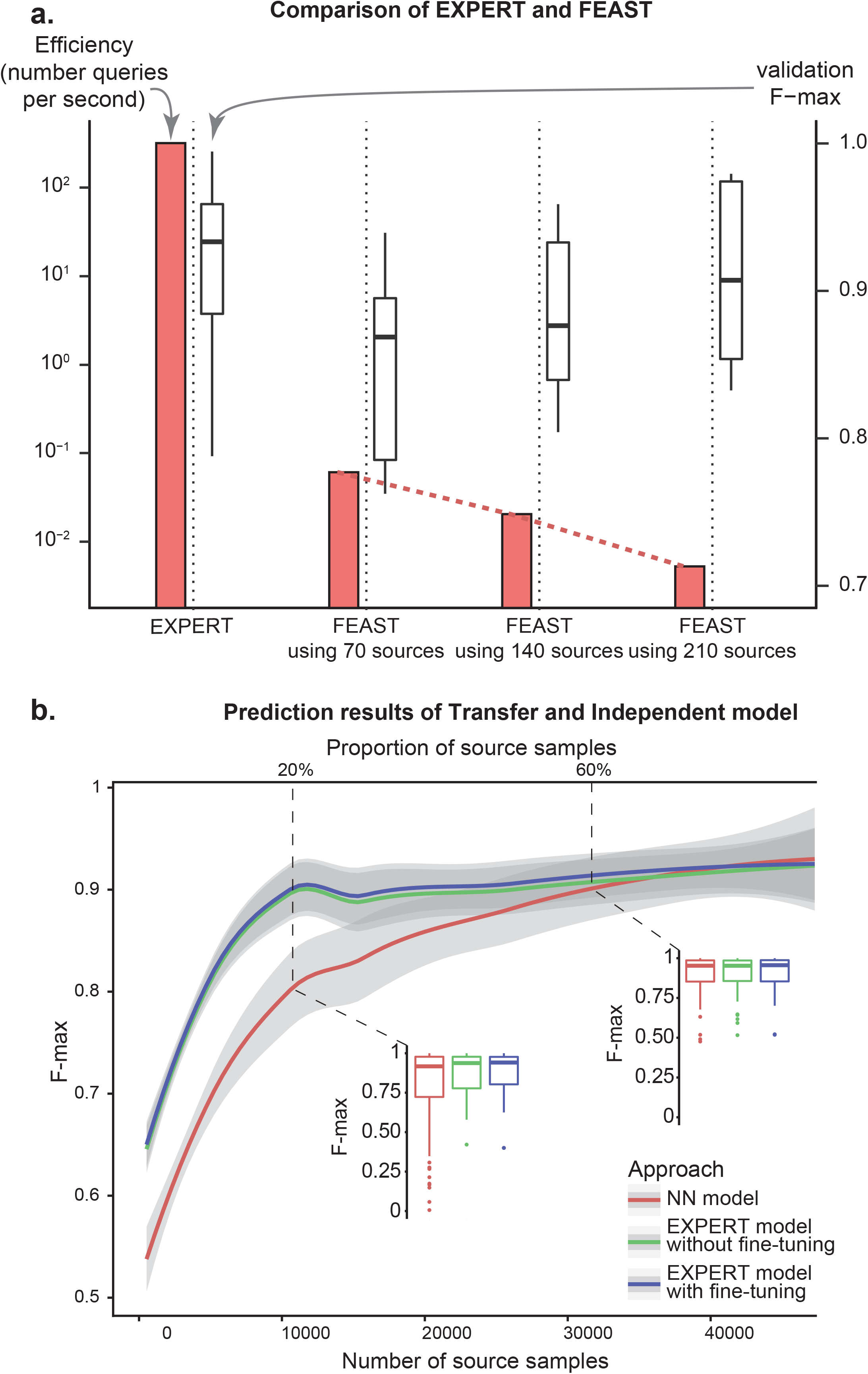
Efficiency, accuracy, and adaptivity of EXPERT. **a**. Comparison of Transfer (GM) EXPERT model with FEAST on efficiency (number of queries/sinks per second, left Y-axis) and accuracy (based on cross-validation, right Y-axis). For FEAST, the sources were randomly selected 70, 140, and 210 samples (10, 20, and 30 samples per biome, respectively). EXPERT’s performance was measured by contextualizing the general model. **b**. The performances (validation F-max, Y-axis) of three models along with different proportions of sources used (X-axis). The NN model was trained solely based on contextual data. The results were obtained by using cross-validation and different proportions (1-10% by a step size of 1%, and 10-90% by a step size of 10%) of source samples. Loess regression was applied to these points using the number of source samples used and F-max.

We have compared the performance of EXPERT with FEAST, as SourceTracker was similarly accurate but slower than FEAST^9^. To compare EXPERT’s accuracy and efficiency with FEAST, we considered community samples from seven biomes (**Supplementary Fig. S3**) as sources, and randomly selected small sets of community samples out of these for comparison as well (**Fig. 2a, Methods**). As a result, EXPERT could simultaneously reach high accuracy and efficiency (Maximal F1-measure F-max = 0.923, over 200 queries/second, **Fig. 2a**). While FEAST faces a severe tradeoff between accuracy and efficiency: FEAST’s accuracy improves as it uses more samples for each biome (F-max = 0.847, 0.884, and 0.911) while efficiency declines nearly exponentially (0.06, 0.02 and 0.005 queries/second, **Fig. 2a, Supplementary Table S6**).

We also compared EXPERT’s accuracy with the NN approach, by using different proportions of source samples (**Fig, 2b**). As the NN approach cannot be directly applied in this context (i.e., NN has no adaptivity), we have manually implemented a model for the comparison. The result showed that the EXPERT model outperforms the NN approach on accuracy: while the MST accuracy steadily increased with the increasing proportion of source samples used, the EXPERT model only required 10% of source samples to achieve a validation F-max of 0.814, while the NN approach required three times as many samples to reach a similar validation F-max of 0.813 (**Supplementary Table S7**). This demonstrated that EXPERT models were able to “understand” the contextualized microbial community profiles based on only a small fraction of samples. Notably, as the fine-tuning optimization clearly improved the accuracy (**Fig. 2b**), the knowledge transfer with fine-tuning was considered the default setting in the following sections.

### Adaptation to newly introduced microbiome data

In this context, we aim to validate EXPERT’s utility in adapting to newly introduced microbial community samples. Such data could be obtained through new sequencing and analytical technologies or originate from rarely studied environments. To test EXPERT’s capability in such context, in addition to the 118,592 communities accessed as of Jan. 2020 from MGnify (referred to as “baseline data”, **Supplementary Table S1, S2, Fig. 3a**), we selected 34,209 communities from MGnify between Jan. 2020 and Oct. 2020 (referred to as “newly introduced data”, **Fig. 3a, Supplementary Table S1, S8, Fig. S4**). Among the newly introduced data, there are 30,788 communities that originated from biomes included in the baseline data as well, and 3,421 communities that originated from newly introduced biomes (**Supplementary Fig. S4**).

**Figure 3.**
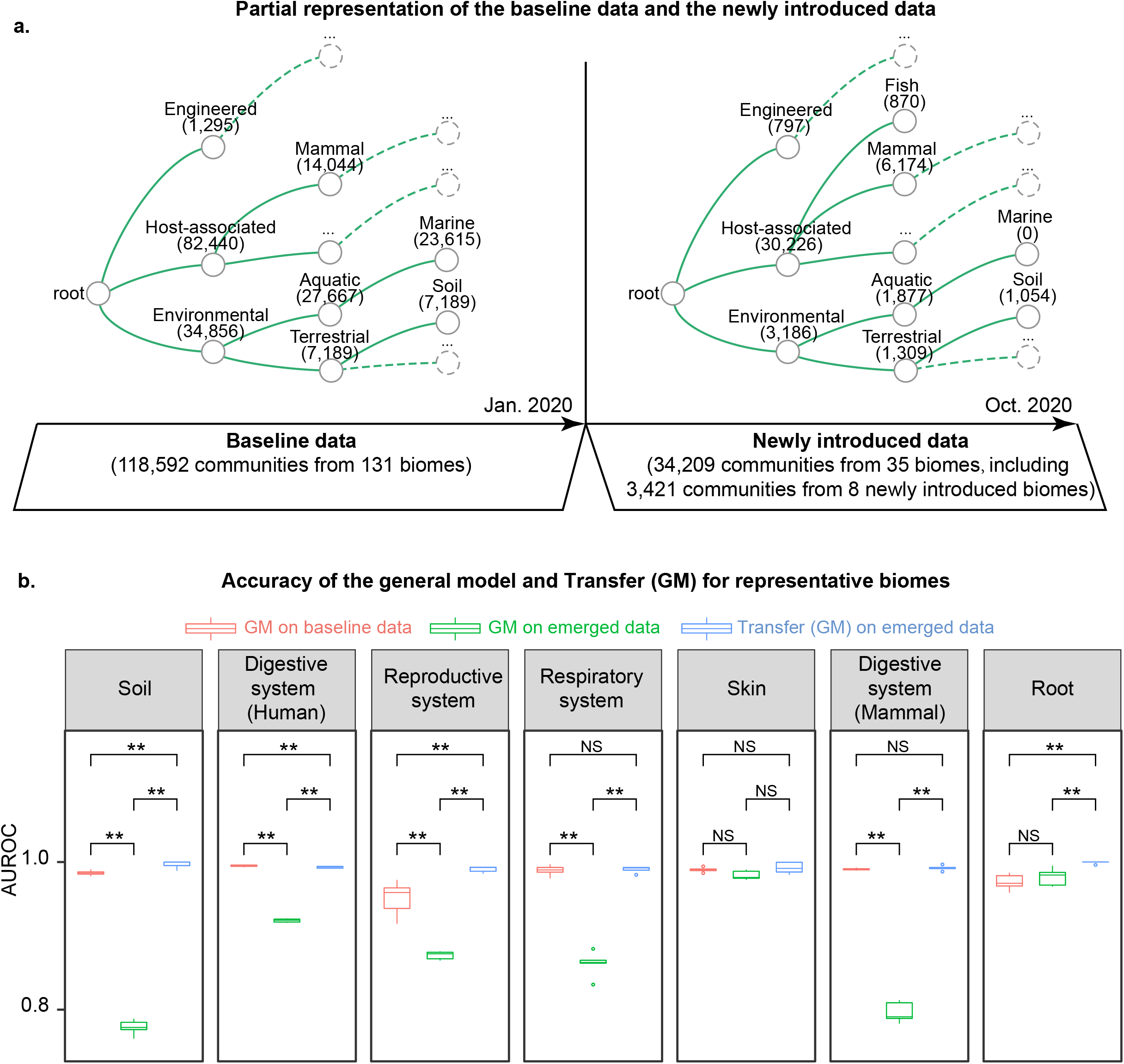
Robust adaptation to the newly introduced microbiome data. **a**. Partial representation of the baseline data and the newly introduced data (with sample size annotations) used to measure the impact of batch effects on MST models and assess the utility of EXPERT. The baseline data contains 118,592 communities deposited before January 2020. The newly introduced data contains 34,209 communities deposited between January 2020 and November 2020, including several newly introduced biomes (e.g. fish-associated biomes). **b**. Performance of EXPERT models on the baseline data and the newly introduced data (performance for seven representative biomes). Furthermore, we can also adapt a fundamental model to newly introduced sources to evaluate these potential microbial sources. Abbreviations: “**”: significant difference; “NS”: non-significant difference; GM: the general model; Transfer (GM): the contextualized model based on the general model. Representative biomes: biomes in the fourth layer of the MGnify biome hierarchy and with sample size greater than 100 in both two datasets.

We first tested the applicability of the general model and EXPERT framework on the newly introduced data. In this context we only considered the 30,788 communities. We directly applied the general model (built based on the baseline data, AUROC = 0.982 by cross-validation) on the data, and obtained a much lower accuracy (AUROC = 0.884, **Supplementary Note 2**). The reason behind this might be the data heterogeneity and batch effect between the two datasets (**Supplementary Fig. S5**). However, by using EXPERT, we could adapt the general model to the newly introduced data, reduce the influence of batch effect on source tracking analysis, and maintain or even further improve the accuracy (AUROC = 0.993, **Fig. 3b**).

We also tested the applicability of EXPERT on the newly introduced biomes, the results have shown that based on the EXPERT framework, the general model could also adapt to the newly introduced biomes (AUROC = 0.988), though the newly introduced biomes were not included in the baseline data. As demonstrated by these results, EXPERT has the potential for extending fundamental models into previously unexplored contexts.

### Context-aware microbial source tracking applications

We then demonstrate EXPERT’s utility in context-aware MST, by focusing on patterns of the human gut microbiome in different contexts: (1) early development of gut microbial communities for infants, (2) association of gut microbial communities with different types of diseases, and (3) association of gut microbial communities with the progression of colorectal cancer. In these contexts, we consider the quantified source contribution generated from EXPERT as a measure to determine the host status.

### The succession of infant gut microbial communities

We next used EXPERT to characterize small compositional changes among infant gut microbial communities during the first year of life. Under this circumstance, we could investigate the dynamic patterns of gut microbial communities from a specific period of life. For instance, if infant samples from multiple time points and sources are present, EXPERT can estimate how much of microbial community in the infant’s gut originated from birth and subsequent time points. To confirm this capability, we used longitudinal data from Backhed et al.^14^, including fecal samples from 98 infants and their mothers, delivered by vaginal delivery or cesarean section (**Fig. 4a, Supplementary Table S1 and S9**). In this part of the study, we considered samples from infants at 12 months of age as queries, and samples from earlier time points or mothers as sources.

**Figure 4.**
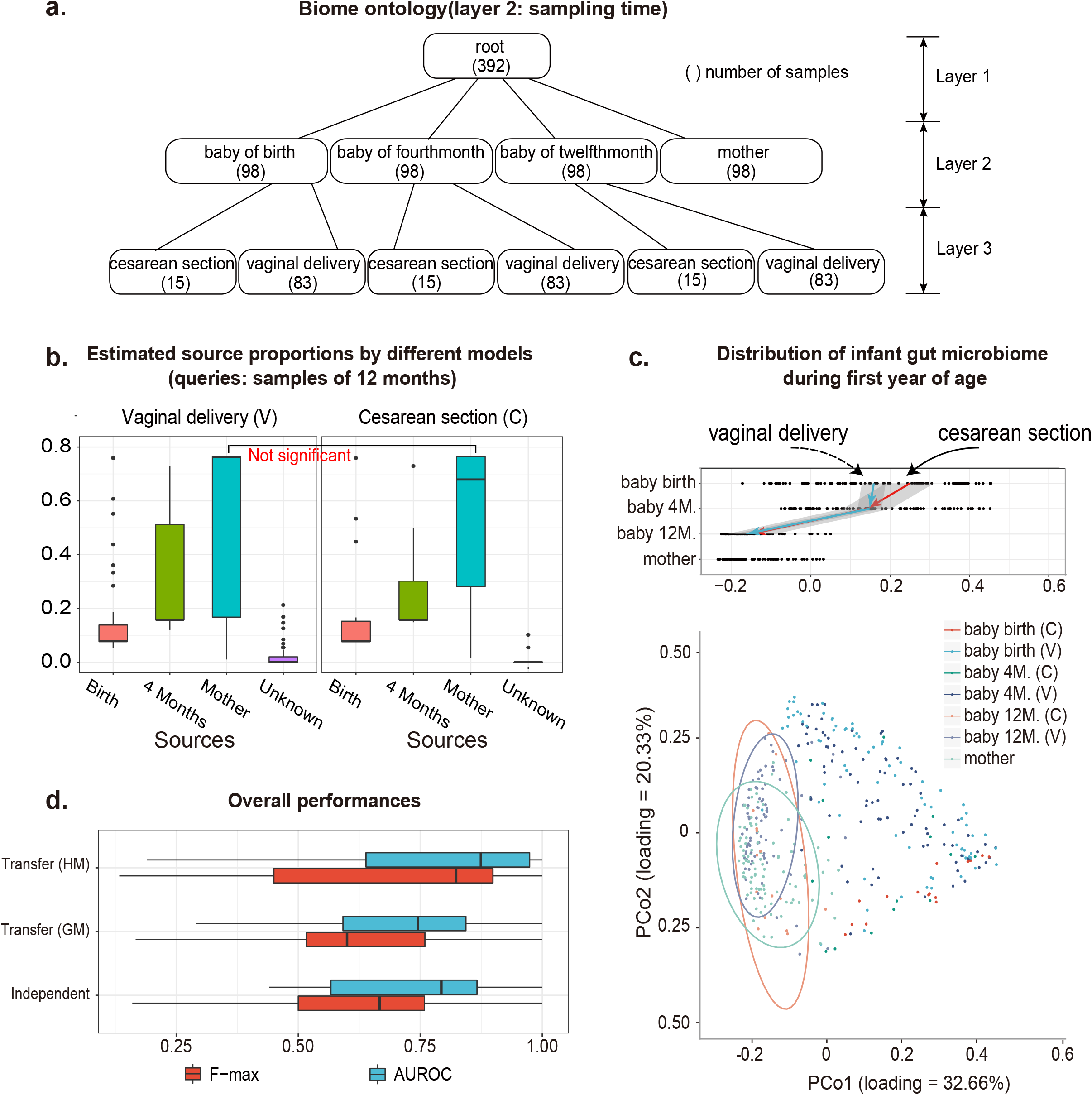
EXPERT’s performance in characterizing gut microbial community development over time for infants. **a**. The hierarchy representing source environments, corresponding to infant samples collected from the ENA database. Environments in the second and third layers were grouped by sampling time and delivery modes. For this part of the study, sources include the gut microbiome of the mother, infant at birth, and four months, queries include the gut microbiome of the infant at 12 months. **b**. Estimated contributions by Transfer (HM) model, separated by two delivery modes. **c**. Distribution of infant gut microbial communities during their first year, using principal-coordinates analysis (PCoA) and distance metric of Jensen Shannon divergence. The dotted line refers to samples delivered vaginally, and the full line refers to samples delivered via cesarean section. The baby of 4 months is abbreviated to baby 4M, the baby of 12 months is abbreviated to baby 12M. The letters “C” and “V” stand for cesarean section and vaginal delivery, respectively. Top panel: samples from the infant’s gut are plotted according to their source and collection date on the Y-axis, and position on the X-axis is plotted according to their first principal coordinate in the PCoA. **d**. The overall performance of models generated based on different fundamental models, in which the Independent model was solely based on the samples used in this context; Transfer (GM) and Transfer (HM) refer to models built based on the general model and human model with fine-tuning, respectively.

Based on the hierarchy that divided samples by sampling time first followed by delivery mode (**Fig. 4a**), we noticed that for infant gut microbial communities at 12 months of age, the maternal contribution is dominant (**Fig. 4b**). Moreover, there is no significant difference in the maternal contribution between cesarean-born and vaginal-born infants (Wilcoxon test, p = 0.929, **Fig. 4b**), consistent with Principal Coordinate Analysis (PCoA) using distance metric either in weighted-UniFrac^15^ or Jensen Shannon divergence^16^ (**Fig. 4c** and **Supplementary Fig. S6**). We concluded that the infant gut at 12 months is largely adapted to exposed environments, resulting in an insignificant difference between samples collected from hosts of different delivery modes, consistent with previous studies^17,18^.

We then assessed the utility of different fundamental models in this context by also introducing the general model and changing the source biome hierarchy (**Supplementary Fig. S7, S8**). We found that the human model can facilitate MST in this context with significantly better performance compared with the general model (Transfer (HM) AUROC = 0.773, Transfer (GM): AUROC = 0.720, Wilcoxon test, p = 0.072), suggesting the use of the human model in this application. Therefore, we suggest that when using EXPERT, it is necessary to choose a proper fundamental model according to the specific context (**Fig. 4d**).

### EXPERT reveals disease-specific patterns within gut microbial communities

The pattern of gut microbial communities could be disease-specific^19^, reflecting the distinct inflammation patterns across diseases. In this context, we aimed to demonstrate EXPERT’s utility in characterizing human gut microbial communities associated with different types of diseases. Using EXPERT, we can measure patterns across multiple diseases. Specifically, we assembled a large gut microbial community dataset, including 13,462 communities representing 19 diseases (**Fig. 5b**) and healthy controls, collected from 101 studies and 27 countries **(Fig. 5a, Supplementary Table S1**, and **S4**). We also introduced the human model to characterize these diseases. By randomly selecting 10% samples of the dataset as queries, and considering the remaining samples as microbial sources, we aim to characterize the pattern across (1) different patients of the same disease, and (2) patients with different diseases. The results revealed that, except for Crohn’s disease, the pattern is shared across patients with the same disease, but not shared across patients with different diseases (**Fig. 5c**). This is consistent with a previous study^19^, which found disease-specific patterns within the human gut microbial communities. Among these diseases examined in this study, we discovered the disease-specific pattern to Liver Cirrhosis and Irritable Bowel Syndrome, which had not been reported in a previous cross-disease study^19^.

**Figure 5.**
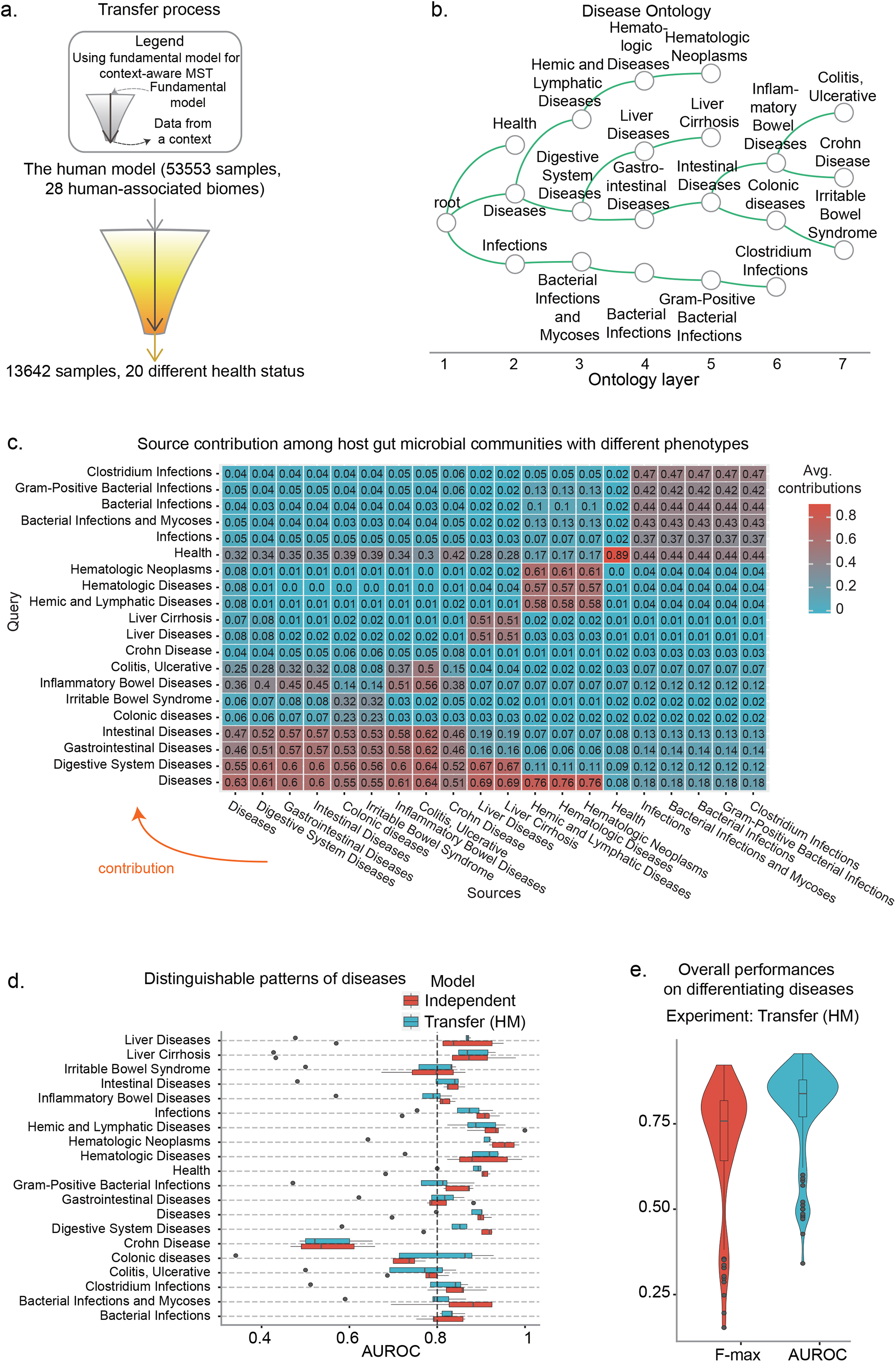
EXPERT reveals disease-specific patterns within gut microbial communities. **a**. Illustration of knowledge transfer utilized for disease pattern analysis. The knowledge transfer between MST contexts was illustrated using different colors (white for human-associated biomes, yellow for gut microbiota-associated disease status). In this analysis, the knowledge from the human model was contextualized (transferred) to the dataset containing 13,642 samples and 19 diseases as well as healthy control. **b**. The hierarchical organization of 19 diseases and healthy control. The hierarchy was constructed by referring disease names to Medical Subject Headings and Human Disease Ontology. The hierarchy includes 20 different health statuses (19 different diseases and infections, plus healthy control) distributed in seven different layers (X-axis). **c**. Average source contribution among all diseases and healthy control, obtained by quantifying contribution from 90% samples (randomly selected) of the dataset to the remaining 10% samples, using Transfer (HM) model based on the human model. The process of random selection and quantification was repeated five times. The heatmap was obtained by averaging the contributions to all samples from each one out of 19 diseases and healthy control. There is no sample overlap between source samples and query samples. **d**. The performance of the EXPERT models on the gut microbial community associated with each disease or healthy control, evaluated based on the source contribution (same as in **Fig. 5c**) and biome-specific evaluation (**Methods**). The dashed line indicates an AUROC of 0.800. **e**. The overall performances of the Transfer (HM) model. Settings of quantification and assessment were the same as **Fig. 5d**.

We further validated the disease-specific patterns by utilizing the Independent model, which was constructed entirely from the same samples. We found that both Independent model and Transfer (HM) model could distinguish diseases with high AUROC, and confirmed that the gut microbial communities may be used to discriminate between these diseases (AUROC over 0.800 for most phenotypes, **Fig. 5d,e**). This demonstrated the utility of EXPERT in large-scale MST analysis, particularly when comparing a wide variety of microbial communities from multiple environments.

### EXPERT characterizes gut microbial communities during cancer progression

Gut microbial communities undergo compositional changes as cancer progresses, and this can be observed in the human gut microbiota, which has been shown to influence the progression of colorectal cancer (CRC)^20^. In this context, we demonstrate EXPERT’s utility in characterizing the progression of CRC using human gut microbiota. We assessed the applicability of EXPERT by leveraging the disease model with Transfer Learning (**Fig. 6a**). We considered 635 samples from five stages in the progression of CRC: 0 (Healthy control) I, II, III, and IV according to the study of Zeller G. et al.^21^ (**Fig. 6b, Supplementary Table S1, and S10**). Preliminary analysis using traditional methods^15,16^ could not show the compositional shifts of the human gut within such progression (**Fig. 6c, Supplementary Fig. S9**). However, by randomly selecting 10% of the dataset as queries, and estimating their resemblant signatures from the remaining samples using the Transfer (DM) model, we found that for gut microbial communities at each CRC stage, a large proportion of microbes could also be found in the communities at the same stage: 0.24, 0.49, 0.56, 0.37 and 0.51 for stage 0 to IV (**Fig. 6d**). These results indicated the association between gut microbiota and CRC progression and suggested the potential of gut microbiota for tracking the progression of CRC^21,22^.

**Figure 6.**
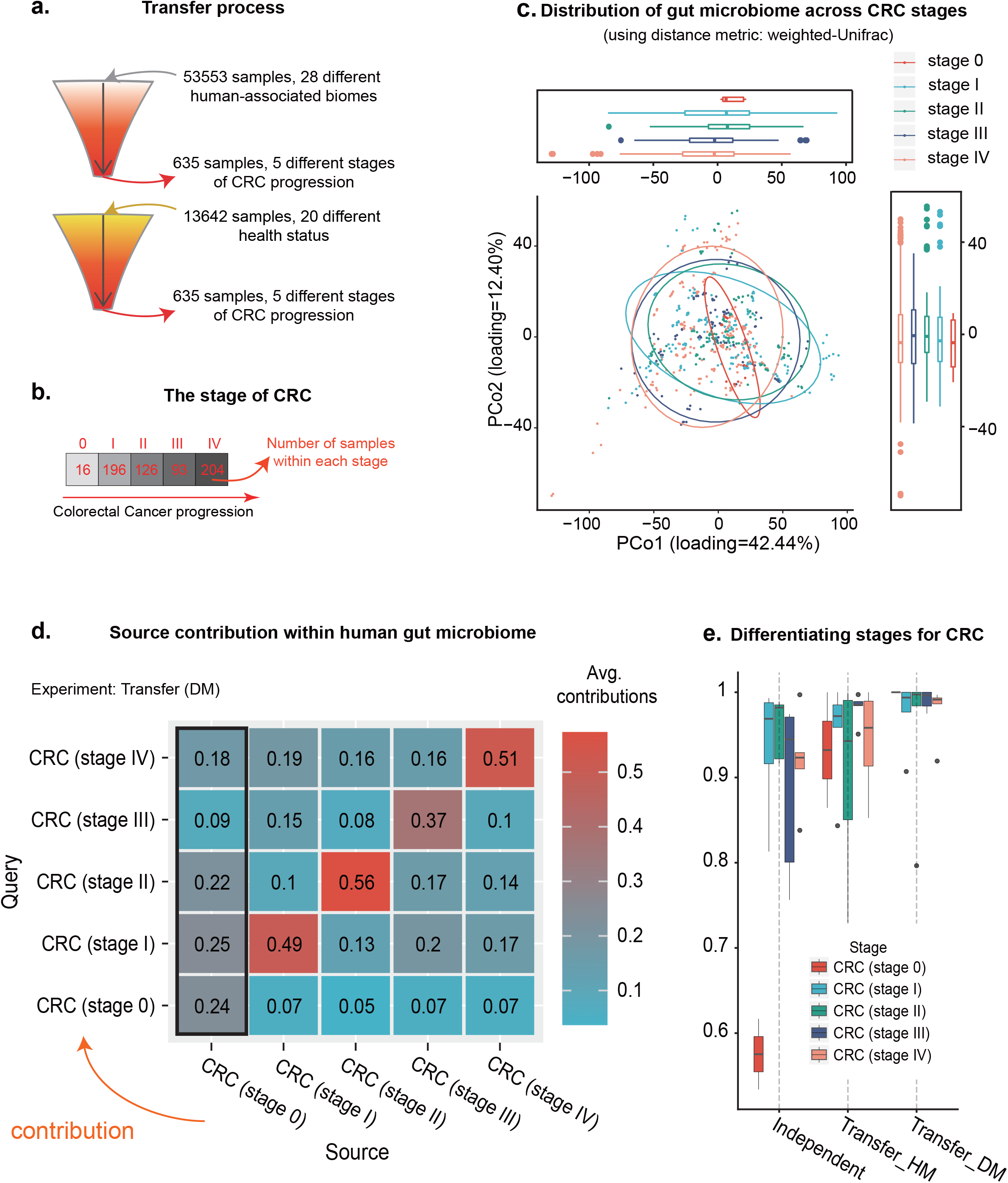
EXPERT characterizes compositional shifts within host gut microbiota during CRC progression. **a**. Illustration of knowledge transfer utilized for characterizing the compositional shifts. The knowledge from the human model learned from 52,537 human-associated communities, as well as the disease model learned from 13,642 human gut communities associated with 19 diseases and healthy control, were transferred to characterize the CRC-related compositional shifts. **b**. The five stages of CRC progression, and the number of samples for each stage. Stage 0: healthy control. **c**. The distribution of gut microbiomes, visualized by PCoA (using distance metric of weighted-UniFrac). **d**. The average contribution of different stages of CRC. The source samples were randomly selected 90% out of the entire dataset. The query samples were the remaining 10% samples. This process of random selection and quantification was repeated five times. There is no sample overlap between source samples and query samples. **e**. The stage-specific performances (AUROC) of EXPERT on different CRC stages (see **Methods** for details of stage-specific evaluation).

We also assessed the EXPERT’s capability in monitoring the progression of CRC, by comparing the performance of different models: For comparison, we generated a Transfer (HM) and an Independent model (solely based on the CRC samples) in addition to the Transfer (DM) model. Results have shown that Transfer (DM) achieved a better performance (AUROC = 0.977, **Fig. 6e, i**) among these three models, highlighting the EXPERT’s utility on tracking the different stages of CRC progression using gut microbial communities.

## Discussion

Broadly, EXPERT adopted a Transfer Learning approach to profoundly elevate and expand the applicability of source tracking, enabling biologically informed novel microbial knowledge discovery. Based on the NN approach and Transfer Learning technique, it could quickly adapt the supervised model for source tracking tasks in different contexts, thus providing a fast, accurate, and context-aware computational approach that enables MST analyses in diverse contexts, for in-depth knowledge discovery.

Our analytical results have confirmed that EXPERT has enabled MST with high speed and fidelity, without the need for pre-defined source samples. Additionally, EXPERT could adapt the fundamental models to newly introduced data, and help reduce the influence of data heterogeneity and batch effects. More importantly, we have shown that MST solely based on the fundamental models may be biased by batch effects, whereas EXPERT can significantly mitigate this influence.

We have demonstrated EXPERT’s utility in context-aware MST in several applications. First, EXPERT can characterize the tiny compositional difference associated with environmental changes. By adapting the human model to microbial communities of infant gut across delivery modes, we found that due to environmental exposure, cesarean-born infants have a largely restored gut microbial community compared with infants born vaginally, consistent with the results of other published analyses^17,18^. Secondly, we demonstrated the utility of EXPERT beyond traditional MST methods by incorporating a dataset of multi-disease gut microbial communities. By using EXPERT on the dataset, we discovered that the human gut microbial community exhibits disease-specific patterns, which is consistent with previous cross-disease research^19^. Thirdly, we showed EXPERT’s utility in characterizing the gut microbiota for patients at various stages of CRC. By using communities from five stages of CRC progression, we found hosts sampled at the same stage shared similar gut microbial communities, enlightening us to realize that the compositional changes within gut microbial communities could reflect the progression of CRC, supported by Shaoming Z et al.^22^.

Several issues need to be looked into further in the future: We noted that in certain contexts (e.g., characterizing gut microbial communities during cancer progression), the accuracy could be improved if the fundamental model was properly selected by referring to the standard ontology^23,24^. EXPERT should provide a collection of fundamental models to enable effective adaptation in diverse MST contexts (e.g., environmental source tracking^28^), and provide an approach for intelligently selecting appropriate fundamental models for a given context. Additionally, the application of EXPERT on the newly introduced data has indicated its robustness against the batch effect, while the extent to which Transfer Learning could overcome the batch effect in microbiome context requires further assessment.

In conclusion, EXPERT enabled accurate and rapid source tracking, as well as biologically informed novel microbial knowledge discovery, by utilizing a Transfer Learning approach. We have demonstrated the applicability of Transfer Learning in the discovery of microbiome knowledge using this method, particularly when dealing with newly introduced data or context-dependent settings. We believed that EXPERT could facilitate high-fidelity source tracking in a broad range of applications.

## Methods

### Datasets

We used six datasets to assess the utility of EXPERT (**Supplementary Table S1**). The hierarchy is essentially a refined subset of an ontology (e.g., Environmental Ontology^23^ or the Human Disease Ontology^24^) or self-defined according to the context of MST. Refer to **Supplementary Note 3**,**4** and **Supplementary Table S11** for the unified data processing pipeline used in the study.

For systematic assessment of our general model, the dataset was obtained from MGnify, which consists of 118,592 communities collected from 131 biomes. Among them, 52,537 samples originated from human biomes, 14,045 samples originated from mammal biomes, 7,189 samples originated from terrestrial biomes, 27,667 samples originated from aquatic biomes. These samples were analyzed by MGnify^25^ before January 2020 (**Supplementary Table S2**). The source environment hierarchy is constructed by referring to the hierarchical biome classification from MGnify database^25^ and the ecosystem classification paths from GOLD database^26^ (**Supplementary Table S12**).

For systematic assessment of our human model, the dataset was a part of the first dataset, in which 52,537 communities from 27 human biomes were selected (**Supplementary Table S3**). The source environment hierarchy is constructed by referring to the hierarchical biome classification from MGnify database^25^ and the ecosystem classification paths from GOLD database^26^.

We also used the newly introduced data in 2020 from MGnify^25^. Which consists of 34,209 communities collected from 35 biomes. Throughout the dataset, 3,421 samples belonging to 8 biomes were newly added by MGnify^25^ after January 2020 (**Supplementary Table S8**). The source environment hierarchy is constructed by referring to the hierarchical biome classification from MGnify database^25^ and the ecosystem classification paths from GOLD database^26^.

For source tracking the succession of infant gut microbiome, the dataset was obtained from MGnify^25^ which consists of 392 fecal samples collected from 98 infants and their biological mothers. Among them, 85 infants were born by vaginal delivery and **13** infants were born by cesarean section. The infant samples were collected at three time points including birth, 4 months, and 12 months. The maternal samples were collected during the first week after delivery (**Supplementary Table S9**).

For disease modeling, the dataset was obtained from GMrepo^27^, including 13,642 communities collected from feces of hosts diagnosed with 19 diseases as well as healthy controls, **Supplementary Table S4**). The source environment hierarchy is constructed by referring to NCBI MeSH^28^ and Human Disease Ontology^24^.

For cancer monitoring, the dataset was obtained from GMrepo^27^, which consists of 16, 93, 126, 196, and 204 communities respectively collected at CRC stage 0, I, II, III, and IV, 635 in total (**Supplementary Table S10**). The source environment hierarchy is constructed by referring to the five stages of CRC.

### The EXPERT framework

#### The EXPERT model

Considering a query sample ***q*** represented by its community structure, as well as its potential sources represented by a hierarchy ***O***, to quantify contributions ***ŷ***_***q***_ from the sources to ***q***, we employed an adaptive and Multi-task NN to learn a mapping ***M*** from a series of source samples ***s* ∈ *D***_***s***_ to their biome sources, 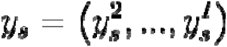 (where 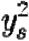 is biome source for source sample in the second layer of the biome hierarchy), and then apply ***M*** on ***q*** to determine the contributions for the query community.

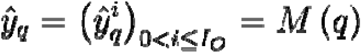

#### Fast inference via forward propagation

We adopt the rationale of Multi-task Learning^13^. EXPERT integrates the representation of each lower layer (which is calculated by its “inter” modules ***M***_***inter***_) into its higher layer, by employing several “integ” modules ***M***_***integ***_. Therefore, together with “output” module ***M***_***output***_, the representation of the contributions is given by

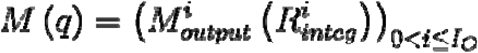

Where

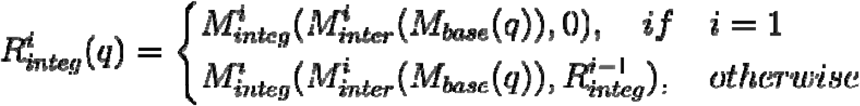

The NN structures of these modules are described in the subsection Adaptive Neural Network.

#### Robust optimization via backward propagation and Transfer Learning

We adopt the rationale of Transfer Learning^12^. Considering 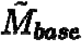 of a fundamental model as a static mapping, the parameters of the rest modules *ŵ* could be solved using gradient descent as well as backpropagation algorithm^29–31^:

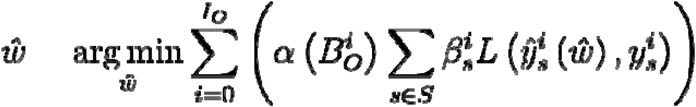

Where

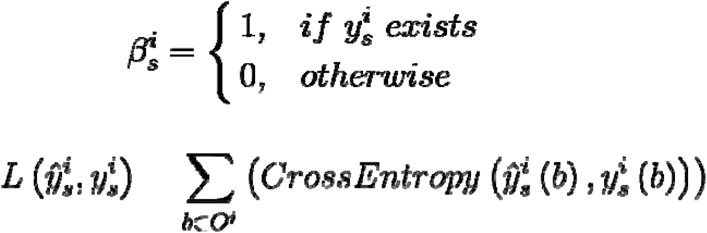

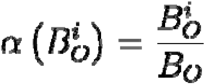 stand for the assigned loss weight for ***i***-th layer (i.e., (***i*** − 1)-th task in the multiple task). 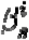 stand for the sample weight assigned for a sample ***s*** on (***i*** − 1)-th task during learning, enabling the learning from partially labeled data. 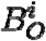 stand for the number of biomes contained in the ***i*** -th layer of the biome hierarchy ***O. O***^***i***^ stand for the ***i***-th layer of the biome hierarchy ***O*.*b* ∈ *O*** ^***i***^ .is a biome in the biome hierarchy ***i***-th layer of the biome hierarchy ***O***.

Then, optimizing the parameters of the entire model (including 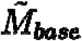), the parameters of the entire model ***w*** can be solved by using gradient descent as well as backpropagation algorithm^29–31^.

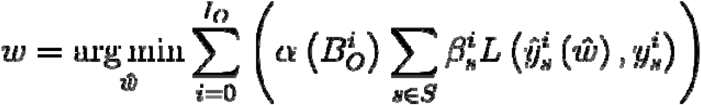

For independent optimization (optimization based on completely random initialization), EXPERT directly optimizes the entire model. See **Supplementary Note 1** for a detailed description for optimization.

#### Adaptive Neural Network

NN approach has limited capability when there is a series of newly introduced source environments, as researchers need to modify the NN model at the code level and re-tune its hyper-parameters. We developed EXPERT’s NN model that changes internal NN structure according to source environments in different contexts, namely the adaptive NN model (**Supplementary Fig. S1**). The EXPERT framework initializes the model according to the hierarchy representing source environments. In the model, there are four conceptual modules.

To extract low-level representations for input data, the model employs the “base” module with two Dense NN layers. The NN layers have fixed structures of 1,024 and 512 neurons, and use ReLU activation^32^ and He initializer with Uniform distribution^33^.

To extract representations that are specific to different hierarchy layers, the model employs the “inter” module with three adaptive Dense NN layers. Denoting ***n*** as the number of source environments in each hierarchy layer, the three NN layers have adaptive structures of, **8 ×*n***, **4×*n***, and **2 ×*n*** neurons, respectively. The three NN layers use ReLU activation^32^ and He initializer with Uniform distribution^33^.

To integrate representation of different hierarchy layers, the model employs the “integ” module with a Concatenation NN layer and an adaptive Dense NN layer. Denoting the number of source environments in each hierarchy layer as ***n***, the NN layer has adaptive structures of **1.5 × *n*** neurons, and uses Tanh activation and Xavier initializer with Uniform distribution^34^.

To estimate according to the integrated representations of different hierarchy layers, the model employs the “output” module with an adaptive Dense NN layer. Denoting the number of source environments in each hierarchy layer as ***n***, the NN layer has adaptive structures of ***n*** neurons, and uses Sigmoid activation and Xavier initializer with Uniform distribution^34^.

### Performance measures

To assess the performance of EXPERT models and other methods, we used these measures:

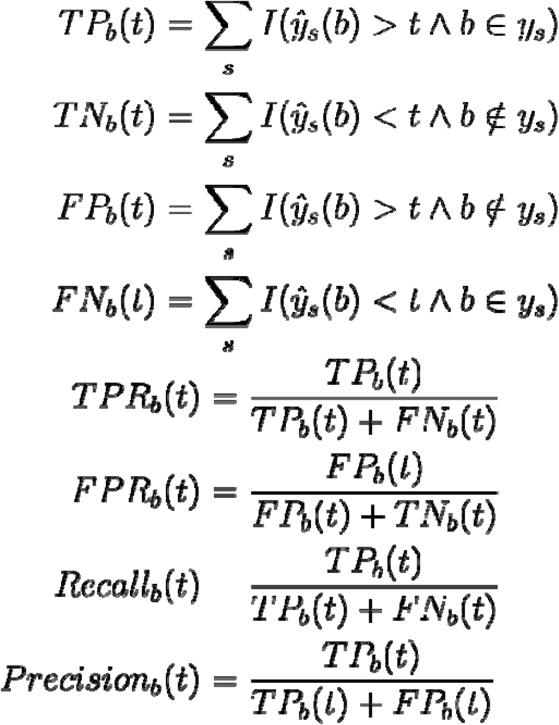

Where ***TP*** is true positive, ***TN*** is true negative, ***FP*** is false positive, ***FN*** is false negative, ***y***_***n***_ **(*b*)** is the quantified contribution from a biome source ***b*** for a microbial community sample ***s***, threshold *t* ∈ [0,1] with a step size of **0.01, *y***_***n***_ is a set of actual biomes for a sample ***s***, and ***I*** is a logical operation function, the value of ***I*** is 1 when the result of logical operation is TRUE, else 0.

Then, two evaluation metrics (F-max, AUROC) were introduced. F-max stands for the maximal F1-measure and was calculated with the following formula. AUROC stands for the area under the ROC (Receiver Operating Characteristics) and was calculated using the composite trapezoidal rule.

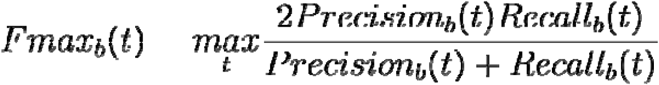

Finally, we treated the average performance across all biomes as the performance of the entire model. Notably, in the subsection “Efficiency, accuracy, and adaptivity of EXPERT”, we only considered biomes with the number of samples > 100 to compute the average performance for the general model, the independent model, Transfer (GM) model, and Transfer (GM0) model.

### Evaluating fundamental models

We assessed each model of the fundamental models through cross-validation, and selected the best model among all trained models as the final model.

We assessed the general model by applying eight-fold cross-validation to the 125,823 microbial community data collected from 132 biomes, and selected the best model among eight trained models as the general model to be transferred.

We assessed the human model by applying repetitive cross-validation (90% as sources to train a model, the resting 10% as queries to test its performance, repeated for five times) to the 52,537 microbial community data collected from 25 biomes, and selected the best model among five trained models as the general model to be transferred.

The assessment of the disease model is the same as the assessment of the human model, but using another dataset consists of 13,462 gut microbial communities associated with 19 diseases.

### Experiment design

We compared EXPERT’s performance with FEAST and the NN approach using the human-associated dataset (**Supplementary Table S1, S3**). We measured the running time using the Linux command “time” and considered the real-time usage for comparison. The efficiency was then calculated using the running time we measured. Refer to **Supplementary Note 5** for detailed comparison procedure for each experiment.

We demonstrated EXPERT’s utility in context-aware MST in three contexts. In these contexts, we used standard hyperparameters for training the model (**Supplementary Note 1**). Detailed descriptions are provided in **Supplementary Note 6**.

### Statistical analysis

Statistical analyses of the contributions have been performed utilizing the Wilcoxon test, at the significance level of□0.05. For all the tests, when the p-value associated is lower than the significance level, one should reject the null hypothesis H0, and accept the alternative hypothesis Ha.

### Visualization of data distribution

Throughout the paper, the box-plot elements are centerline, median; box limits, upper and lower quartiles; whiskers, 1.5 ×□interquartile range (IQR); points, and outliers. The Violin plot is also used for data distribution analysis, mainly for comparison. The PCoA is also used for data distribution analysis, with ellipses representing a confidential interval of 0.95. The Principle Coordination is obtained through applying beta diversity measurement (Scikit-bio version 0.5.6, **Supplementary Table S5**) on the abundance of all taxa in seven ranks, namely Superkingdom, Kingdom, Phylum, Class, Order, Family, Genus, and Species. The source code of the PCoA analysis is hosted on GitHub at https://github.com/AdeBC/UniPCoA.

## Data availability

The collected samples from MGnify and GMrepo databases were annotated with their associated biomes/phenotypes in **Supplementary Table S2-S4, S8-S10**. All the processed data are uploaded and hosted at https://github.com/HUST-NingKang-Lab/EXPERT-use-cases.

## Code availability

All source codes have been uploaded to the website at https://github.com/HUST-NingKang-Lab/EXPERT. Detailed software and models used in this study are provided in **Supplementary Table S5**.

## Acknowledgments

This work was partially supported by National Natural Science Foundation of China grant 32071465, 31871334, and 31671374, and the China Ministry of Science and Technology’s National Key R&D Program grant (No. 2018YFC0910502).

## Author contributions

HC and KN designed the study, conceived of and proposed the idea, designed and developed the framework. HC, QY, GX, NW, SH and XH performed the experiments and analyzed the data. HC, QY, YZ, GX, and NW visualized the data. CS and SW provided valuable data. HC, QY, YZ, WC, LPC, and KN contributed to editing and proofreading the manuscript. All authors read and approved the final manuscript.

## Ethics approval and consent to participate

Not applicable.

