## Supplementary Information for "EXPERT: Transfer Learning-enabled context-aware microbial source tracking"

**Supplementary Figure 1-9, Supplementary Table S1-12 and Supplementary Note 1-6**

### Supplementary Figure 1

**Supplementary Figure 1. Neural network architecture of EXPERT fundamental models.** EXPERT enabled adaptation to new biome ontology by employing a structurally adaptive fundamental model. The model could reinitialize context-dependent layers according to different biome ontology. Among the entire model, two fully-connected layers were utilized to extract low-level representations of input communities. And contextual NN layers were utilized to estimate contribution to the input communities.

### Supplementary Figure 2

**Supplementary Figure 2. Diagram of microbial source tracking process according to hierarchically organized source environments.** The process is exemplified using a microbial community from “coastal” biome and an MST model built based on a hierarchy of a series of human and environmental sources.

### Supplementary Figure 3

**Supplementary Figure 3. Hierarchy representing 27 human-associated biomes as well as their corresponding sample size.** Biomes for comparison with FEAST were shown in the yellow box. Only biomes with the number of samples > 100 are used for evaluation.

### Supplementary Figure 4

**Supplementary Figure 4. Hierarchy representing 35 biomes in the newly introduced data, as well as their corresponding sample size.**

### Supplementary Figure 5

**Supplementary Figure 5. Batch effects of community data between the baseline data and the newly introduced data.** The first principal coordination (PCo1, X-axis) and the second principal coordination (PCo2, Y-axis) were plotted based on PCoA analysis of Scikit-bio (version 0.5.6). The Ellipsoids stand for a confidential interval of 95%.

### Supplementary Figure 6

**Supplementary Figure 6. Distribution of infant gut microbiome during first year of age, using principal-coordinates analysis (PCoA) and distance metric of weighted-Unifrac.** From PCoA, there is no significant difference between infant gut microbiome at 12 months of age of different delivery modes. The first principal coordination (PCo1, with 51.67% variance explained, X-axis) and the second principal coordination (PCo2, with 15.17% variance explained, Y-axis) were plotted based on PCoA analysis of Scikit-bio (version 0.5.6). Baby of 4th month is abbreviated to baby 4M., baby of 12th month is abbreviated to baby 12M. "C" represents cesarean section, "V" represents vaginal delivery.

### Supplementary Figure 7

**Supplementary Figure 7. Estimated source contributions by different models of EXPERT.** We noted that regardless of delivery modes, when source tracking infant gut microbiome at birth, contribution of infant gut at 4th month is dominant. Separated by two delivery modes and transferred from general model (above) and human model (below), where sources include gut microbiome of mother, infant at 4 months and 12 months, queries include gut microbiome of infant at birth.

### Supplementary Figure 8

**Supplementary Figure 8. a.** The ontology structure of biomes, corresponding to infant data from the ENA database. Layer 2 based on delivery modes, while layer 3 based on sampling time. **b.** The overall performance of models generated based on different training schemes. Independent model was based on independent training, the Transfer (GM) model was generated based on the general model, and the Transfer (HM) model was generated based on the human model. The Transfer (HM) model showed the best performance when source tracking the samples. Red represents AUROC, blue represents F-max.

### Supplementary Figure 9

**Supplementary Figure 9. Distribution of gut microbiome across CRC stages, using PCoA and distance metric of Jensen Shannon divergence.** The first principal coordination (PCo1, with 17.43% variance explained, X-axis) and the second principal coordination (PCo2, with 12.32% variance explained, Y-axis) were plotted based on PCoA analysis of Scikit-bio (version 0.5.6). The Ellipsoids stand for a confidential interval of 95%. Note that the analysis was based on abundances of all taxonomies (including each involved taxon from seven ranks: super-kingdom, kingdom, phylum, class, order, family, genus). For code used to generate the PCoA result, please refer to our GitHub (https://github.com/HUST-NingKang-Lab/EXPERT).

### Supplementary Table S1

**Supplementary Table S1. Information about the datasets used in this study.**

| **Description** | **Size** | **Associated metadata** | **Details** | **Source** |
| --- | --- | --- | --- | --- |
| Dataset for systematic assessment of general model | 118,592 | 132 biomes | **Supplementary Table S2** | MGnify database^1^ |
| Dataset for systematic assessment of our human model | 52,537 | 25 biomes | **Supplementary Table S3** | MGnify database^1^ |
| Dataset for disease pattern analysis | 13,642 | 20 disease status | **Supplementary Table S4** | GMrepo database^2^ |
| Dataset for assessing the generalizing performance on newly introduced data | 34,209 | 35 biomes | **Supplementary Table S8** | MGnify database^1^ |
| Dataset for source tracking the succession of infant gut microbiome | 392 | 11 biomes | **Supplementary Table S9** | MGnify database^1^ |
| Dataset for CRC analysis | 635 | 5 stages of CRC | **Supplementary Table S10** | GMrepo database^2^ |

### Supplementary Table S2

**Supplementary Table S2. Associated metadata for assessing the generalization performance in a general setting.** In the table, the run accession in ENA and MGnify databases and associated biome (in total 132) of 118,592 community samples were provided (attached in Excel table). The dataset was introduced to train and assess the general model in the study.

### Supplementary Table S3

**Supplementary Table S3. Associated metadata for assessing the generalization performance in context-dependent settings.** In the table, the run accession in ENA and MGnify databases and associated biome (in total 27) of 52,537 community samples were provided (attached in Excel table). The dataset was introduced to train and assess the human model, and to assess the utility of transfer learning in the study.

### Supplementary Table S4

**Supplementary Table S4. Associated metadata for systematic disease diagnosis.** In the table, the run accession in GMrepo database and associated host phenotypes (in total 20) of 13,642 community samples were provided (attached in Excel table). The dataset was introduced to train and assess the disease model.

### Supplementary Table S5

**Supplementary Table S5. Databases, packages, and models used in our study.**

**Database**

| **Database** | **Ref** | **Website** |
| --- | --- | --- |
| EBI MGnify (accessed as of Nov, 4th, 2020) | Mitchell AL, Almeida A, et al. 2020 | https://www.ebi.ac.uk/metagenomics/ |
| EBI ENA (accessed as of Nov, 4th, 2020) | Rasko Leinonen, Ruth Akhtar, et al. 2011 | https://www.ebi.ac.uk/ena/browser/home |
| NCBI MeSH (accessed as of Sep, 30th, 2020) | Eric W Sayers, Jeff Beck, et al. 2019 | https://www.ncbi.nlm.nih.gov/mesh/ |
| NCBI taxdump (released Sep 1st, 2020) | Eric W Sayers, Jeff Beck, et al. 2019 | https://ftp.ncbi.nlm.nih.gov/pub/taxonomy/taxdump_archive/ |
| GMrepo database (Released Nov 4th, 2020) | Sicheng Wu, Chuqing Sun, et al. 2020 | https://gmrepo.humangut.info/home |

**Software**

| **Software** | **Author** | **Website** |
| --- | --- | --- |
| EXPERT (version 0.2) | Hui Chong | https://github.com/HUST-NingKang-Lab/EXPERT |
| FEAST (committed Aug 30, 2019) | Shenhav, L., Thompson, M. et al., 2019 | https://github.com/cozygene/FEAST |
| Python (version 3.8.2) | Oliphant and Travis E, 2007 | https://www.python.org |
| SQLite3 | Gerhard Haring | https://docs.python.org/3/library/sqlite3.html |
| ETE toolkit | Jaime Huerta-Cepas et al., 2016 | http://etetoolkit.org/ |
| Pandas (version 1.0.1) | McKinney, W. et al., 2010 | https://pandas.pydata.org |
| Treelib (version 1.5.5) | Xiaming Chen | https://github.com/caesar0301/treelib |
| NumPy (version 1.18.5) | Charles R. Harris, K. et al., 2020 | https://numpy.org |
| Scikit-learn (version 0.23.2) | Pedregosa et al., 2011 | https://scikit-learn.org |
| Scikit-bio (version 0.5.6) | The scikit-bio development team, 2020 | http://scikit-bio.org/docs/latest/index.html# |
| SciPy (version 1.5.2) | Pauli Virtanen, Ralf Gommers et al., 2020 | https://www.scipy.org/ |
| TensorFlow (version 2.3.1) | Martín Abadi et al., 2015 | https://www.tensorflow.org |
| R (version 3.6.1) | Team, R. C., 2013 | https://www.r-project.org |
| ggplot2 (version 3.3.3) | Hadley Wickham, 2016 | https://ggplot2.tidyverse.org/index.html |
| Plotnine (version 0.7.1) | Hassan Kibirige et al., 2020 | https://plotnine.readthedocs.io/en/stable/index.html |

**Model**

| **Model** | **Ontology** | **Link** |
| --- | --- | --- |
| General model (GM) | Biome ontology from MGnify (Sep. 2019) | https://github.com/HUST-NingKang-Lab/EXPERT/releases/download/v0.2-m/general_model.tgz |
| Human model (HM) | Ontology for human biomes from MGnify (Sep. 2019) | https://github.com/HUST-NingKang-Lab/EXPERT/releases/download/v0.2-m/human_model.tgz |
| Disease model (DM) | Ontology for human diseases from MeSH (Nov. 2020) | https://github.com/HUST-NingKang-Lab/EXPERT/releases/download/v0.2-m/disease_model.tgz |

### Supplementary Table S6

**Supplementary Table S6. Comparison of Transfer (GM) and FEAST in source tracking accuracy and efficiency.** Query time was extrapolated to 1,000 query samples for easy comparison.

| **Model** | **Source setting** | **Total training time (s)** | **Query time per 1,000 query samples (s)** | **F-max** |
| --- | --- | --- | --- | --- |
| FEAST | 70 source samples  (10 * 7) | - | 16,532 | 0.847 |
|  | 140 source samples  (20 * 7) | - | 48,930 | 0.884 |
|  | 210 source samples  (30 * 7) | - | 190,841 | 0.911 |
| Transfer (GM) | 35,975 source samples | 1,340 | 3 | 0.923 |

### Supplementary Table S7

**Supplementary Table S7. The performance (F-max) of three models across different proportions of the human dataset considered as sources.** The assessment results of Independent, Transfer (GM), and Transfer (GM0) models when different proportions of the human dataset were used for training.

| **Proportion** | **Independent** | **Transfer (GM0)** | **Transfer (GM)** |
| --- | --- | --- | --- |
| 0.01 | 0.503 | 0.58 | 0.581 |
| 0.02 | 0.552 | 0.657 | 0.659 |
| 0.03 | 0.607 | 0.703 | 0.711 |
| 0.04 | 0.589 | 0.738 | 0.736 |
| 0.05 | 0.628 | 0.76 | 0.771 |
| 0.06 | 0.633 | 0.764 | 0.791 |
| 0.07 | 0.666 | 0.789 | 0.79 |
| 0.08 | 0.693 | 0.811 | 0.79 |
| 0.09 | 0.716 | 0.816 | 0.827 |
| **0.1** | **0.704** | **0.81** | **0.814** |
| 0.2 | 0.789 | 0.872 | 0.877 |
| **0.3** | **0.813** | **0.883** | **0.888** |
| 0.4 | 0.882 | 0.89 | 0.898 |
| 0.5 | 0.861 | 0.896 | 0.901 |
| 0.6 | 0.905 | 0.91 | 0.914 |
| 0.7 | 0.91 | 0.912 | 0.919 |
| 0.8 | 0.923 | 0.917 | 0.927 |
| 0.9 | 0.931 | 0.925 | 0.923 |

### Supplementary Table S8

**Supplementary Table S8. Associated metadata for assessing the generalization performance on the newly introduced data.** In the table, the run accession in ENA and MGnify databases and associated biome (in total 35) of 34,209 community samples were provided (attached in Excel table). The dataset was introduced to train and assess the generalization performance of the general model on the newly introduced data, and used in the subsection “Adaptation to newly introduced microbiome data” of the study.

### Supplementary Table S9

**Supplementary Table S9. Associated metadata for source tracking the succession in the infant gut microbiome.** In the table, the run accession in ENA and MGnify databases and associated metadata of 392 community samples were provided (attached in Excel table). The dataset was used in the subsection “The succession of infant gut microbial communities” of the study.

### Supplementary Table S10

**Supplementary Table S10. Associated metadata for monitoring the progression of colorectal cancer.** In the table, the run accession in GMrepo database and associated metadata of 635 community samples were provided (attached in Excel table). The dataset was used in the subsection “EXPERT characterizes gut microbial communities during cancer progression” of the study.

### Supplementary Table S11

**Supplementary Table S11. The phylogenetic tree used in EXPERT.** The tree contains 6006 genera and three taxon ranks (phylum, order, and genus) used for encoding 18018 features for EXPERT (attached in Excel table).

### Supplementary Table S12

**Supplementary Table S12. Representation for general biome ontology.** In the table, each row represents a path from root node to a leaf node and each column represents a layer of the ontology.

| **Layer 1** | **Layer 2** | **Layer 3** | **Layer 4** | **Layer 5** | **Layer 6** |
| --- | --- | --- | --- | --- | --- |
| root | Engineered | Wastewater | Activated Sludge |  |  |
| root | Engineered | Wastewater | Industrial wastewater | Agricultural wastewater |  |
| root | Engineered | Wastewater | Industrial wastewater | Mine water |  |
| root | Engineered | Wastewater | Industrial wastewater | Petrochemical |  |
| root | Engineered | Wastewater | Nutrient removal | Dissolved organics (aerobic) |  |
| root | Engineered | Wastewater | Nutrient removal | Dissolved organics (anaerobic) |  |
| root | Engineered | Wastewater | Water and sludge |  |  |
| root | Environmental | Aquatic | Freshwater | Groundwater | Biofilm |
| root | Environmental | Aquatic | Freshwater | Lake |  |
| root | Environmental | Aquatic | Marine | Coastal | Sediment |
| root | Environmental | Aquatic | Marine | Cold seeps | Sediment |
| root | Environmental | Aquatic | Marine | Hydrothermal vents | Black smokers |
| root | Environmental | Aquatic | Marine | Hydrothermal vents | Diffuse flow |
| root | Environmental | Aquatic | Marine | Hydrothermal vents | Microbial mats |
| root | Environmental | Aquatic | Marine | Hydrothermal vents | Sediment |
| root | Environmental | Aquatic | Marine | Intertidal zone | Coral reef |
| root | Environmental | Aquatic | Marine | Intertidal zone | Estuary |
| root | Environmental | Aquatic | Marine | Intertidal zone | Mangrove swamp |
| root | Environmental | Aquatic | Marine | Intertidal zone | Microbialites |
| root | Environmental | Aquatic | Marine | Intertidal zone | Salt marsh |
| root | Environmental | Aquatic | Marine | Intertidal zone | Sediment |
| root | Environmental | Aquatic | Marine | Marginal Sea |  |
| root | Environmental | Aquatic | Marine | Neritic zone | Sediment |
| root | Environmental | Aquatic | Marine | Oceanic | Abyssal plane |
| root | Environmental | Aquatic | Marine | Oceanic | Aphotic zone |
| root | Environmental | Aquatic | Marine | Oceanic | Benthic |
| root | Environmental | Aquatic | Marine | Oceanic | Oil-contaminated |
| root | Environmental | Aquatic | Marine | Oceanic | Oil-contaminated sediments |
| root | Environmental | Aquatic | Marine | Oceanic | Photic zone |
| root | Environmental | Aquatic | Marine | Oceanic | Sediment |
| root | Environmental | Aquatic | Marine | Oil-contaminated sediment |  |
| root | Environmental | Aquatic | Marine | Oil seeps |  |
| root | Environmental | Aquatic | Marine | Pelagic |  |
| root | Environmental | Aquatic | Marine | Sediment |  |
| root | Environmental | Aquatic | Marine | Volcanic |  |
| root | Environmental | Aquatic | Marine | Wetlands | Sediment |
| root | Environmental | Aquatic | Non-marine Saline and Alkaline | Alkaline | Sediment |
| root | Environmental | Aquatic | Non-marine Saline and Alkaline | Hypersaline | Sediment |
| root | Environmental | Aquatic | Non-marine Saline and Alkaline | Salt crystallizer pond |  |
| root | Environmental | Aquatic | Sediment |  |  |
| root | Environmental | Aquatic | Thermal springs | Sediment |  |
| root | Environmental | Terrestrial | Soil | Agricultural |  |
| root | Environmental | Terrestrial | Soil | Boreal forest |  |
| root | Environmental | Terrestrial | Soil | Contaminated |  |
| root | Environmental | Terrestrial | Soil | Crop | Agricultural land |
| root | Environmental | Terrestrial | Soil | Desert |  |
| root | Environmental | Terrestrial | Soil | Forest soil |  |
| root | Environmental | Terrestrial | Soil | Grasslands |  |
| root | Environmental | Terrestrial | Soil | Loam | Agricultural |
| root | Environmental | Terrestrial | Soil | Permafrost |  |
| root | Environmental | Terrestrial | Soil | Sand |  |
| root | Environmental | Terrestrial | Soil | Tropical rainforest |  |
| root | Environmental | Terrestrial | Soil | Uranium contaminated |  |
| root | Environmental | Terrestrial | Soil | Wetlands |  |
| root | Host-associated | Human | Circulatory system | Blood |  |
| root | Host-associated | Human | Digestive system | Intestine |  |
| root | Host-associated | Human | Digestive system | Large intestine | Fecal |
| root | Host-associated | Human | Digestive system | Oral | buccal mucosa |
| root | Host-associated | Human | Digestive system | Oral | Periodontal pockets |
| root | Host-associated | Human | Digestive system | Oral | Saliva |
| root | Host-associated | Human | Digestive system | Oral | Subgingival plaque |
| root | Host-associated | Human | Digestive system | Oral | Supragingival plaque |
| root | Host-associated | Human | Digestive system | Oral | Throat |
| root | Host-associated | Human | Digestive system | Oral | tongue dorsum |
| root | Host-associated | Human | Lympathic system | Lymph nodes |  |
| root | Host-associated | Human | Reproductive system | Female |  |
| root | Host-associated | Human | Reproductive system | Vagina | posterior fornix |
| root | Host-associated | Human | Respiratory system | Nasopharyngeal | Nasal cavity |
| root | Host-associated | Human | Respiratory system | Nasopharyngeal | Pharynx |
| root | Host-associated | Human | Respiratory system | Pulmonary system | Sputum |
| root | Host-associated | Human | Skin |  |  |
| root | Host-associated | Insecta | Digestive system |  |  |
| root | Host-associated | Mammals | Digestive system | Fecal |  |
| root | Host-associated | Mammals | Digestive system | Foregut | Rumen |
| root | Host-associated | Mammals | Digestive system | Large intestine | Cecum |
| root | Host-associated | Mammals | Digestive system | Large intestine | Fecal |
| root | Host-associated | Mammals | Digestive system | Oral cavity | Buccal mucosa |
| root | Host-associated | Mammals | Digestive system | Stomach | Rumen |
| root | Host-associated | Plants | Phylloplane | Endophytes |  |
| root | Host-associated | Plants | Rhizoplane | Endophytes |  |
| root | Host-associated | Plants | Rhizosphere | Epiphytes |  |
| root | Host-associated | Plants | Rhizosphere | Forest soil |  |
| root | Host-associated | Plants | Rhizosphere | Soil |  |
| root | Host-associated | Plants | Root |  |  |

### Supplementary Note 1

**Supplementary Note 1. Settings for random initialization, independent optimization, and transfer learning**

We used Xavier initialization^4,5^ with uniform distribution for layers with tanh or sigmoid activation, and He initialization^4,6^ with uniform distribution for layers with ReLU activation, following the suggestions in their original studies (**Supplementary Fig. 1**).

We used independent optimization to generate EXPERT models only based on knowledge and data of the context. Firstly, the model was trained using Adam optimizer^7^ with a learning rate of 1x10-5 and a mini-batch size of 1024, for two epochs. Then, the model was trained using Adam optimizer with an initial learning rate of 1x10-3 and a mini-batch size of 1024. The learning rate was divided by a factor of 10 when the test loss stopped optimizing for 5 epochs. And the training was early-stopped when the test loss stopped optimizing for 15 epochs. Before each epoch of optimization, 10% out of training data were randomly selected to validate the model after the epoch. Additionally, the minimum learning rate and maximal epochs of training were set to 1x10-5 and 1000, respectively.

We used transfer learning to generate EXPERT models based on the knowledge of a fundamental model, and data of a context. After weight reuse, the model was trained using Adam optimizer with an initial learning rate of 1x10-3 and a mini-batch size of 64. The same settings of the learning rate decay and early-stopping as independent optimization were used here. Then, the model was fine-tuned using Adam optimizer with a learning rate of 1x10-5, with the same mini-batch size of 64.

### Supplementary Note 2

**Supplementary Note 2. Adaptation to newly introduced microbiome data**

To compare the performance of the general model on different datasets, we first assessed the performance of the general model on the baseline data using eight-fold cross-validation. Then, we repeated a random cross-validation process. For each time of the cross-validation, we considered 90% of the dataset as sources, and the remaining samples as queries. For the general model’s performance on the newly introduced data, we directly searched queries from each time of the cross-validation against the general model to assess its performance. For the Transfer (GM) model’s performance, we re-adapted the general model using the sources from each time of the cross-validation, and evaluated the performance using the queries from each time of the cross-validation. The queries and sources used for assessing the performance of the general model and the Transfer (GM) model are the same. We enabled “fine-tuning” and “update statistics” options when optimizing the Transfer (GM) model. For other hyper-parameters used for training, please refer to Supplementary Note 2 for a detailed description.

### Supplementary Note 3

**Supplementary Note 3. Data preprocessing**

#### *Relative abundance calculation according to reference database*

In order to bypass the impact from the sequencing depth of the rich-sourced data, we firstly regularized the abundance data by calculating relative abundances according to only the taxa mapped to our phylogenetic tree (**Supplementary Table S11**), which is a part of taxonomical classification tree in NCBI taxonomy database^33^ and reflects the input feature set for EXPERT model (as explained below).

#### *Universal feature set for homogeneous Transfer Learning*

In order to realize knowledge transfer among multi-faceted microbial source tracking applications, we utilized a uniform collection of features in all these applications. Such a universal feature set was established according to only the variance of relative abundances at genes level: Among the first dataset (for which the most comprehensive), 6,006 genera with variance of relative abundances above the threshold of 10^-3^ times the average variance across all genera. We constructed a phylogenetic tree based on the classification of these 6,006 genera (three taxon ranks were included: “phylum”, “order”, “genus”), and served the tree for abundance mapping (**Supplementary Note 1, Supplementary Table S11**).

#### *Z-score standardization of relative abundances*

In order to speed up the optimization of EXPERT model, we standardized the relative abundances before feeding them into the model, by applying z-score standardization on normalized relative abundances.

### Supplementary Note 4

**Supplementary Note 4. Detailed data preprocessing procedure and parallelized abundance aggregation.**

The community data generated by a variety of analysis pipelines often have different forms of taxa representation. To feed them to EXPERT, we retrieved their classification lineages from NCBI taxonomy database, remapped their abundances to our phylogenetic tree (a part of NCBI taxonomy tree), and recalculated the relative abundances according to only those successfully mapped taxa.

The EXPERT model accepts abundances in genus level as inputs, and calculates abundances in phylum and order levels automatically. This has been achieved in a rapid manner: EXPERT utilizes matrix operations of TensorFlow^4^ to speed up such processes, rather than traverse and add up the abundance over the phylogenetic tree from the bottom-up. Specifically, according to taxa classification information of the phylogenetic tree, we first calculated two abundance aggregation matrices for two higher levels "phylum" and "order":

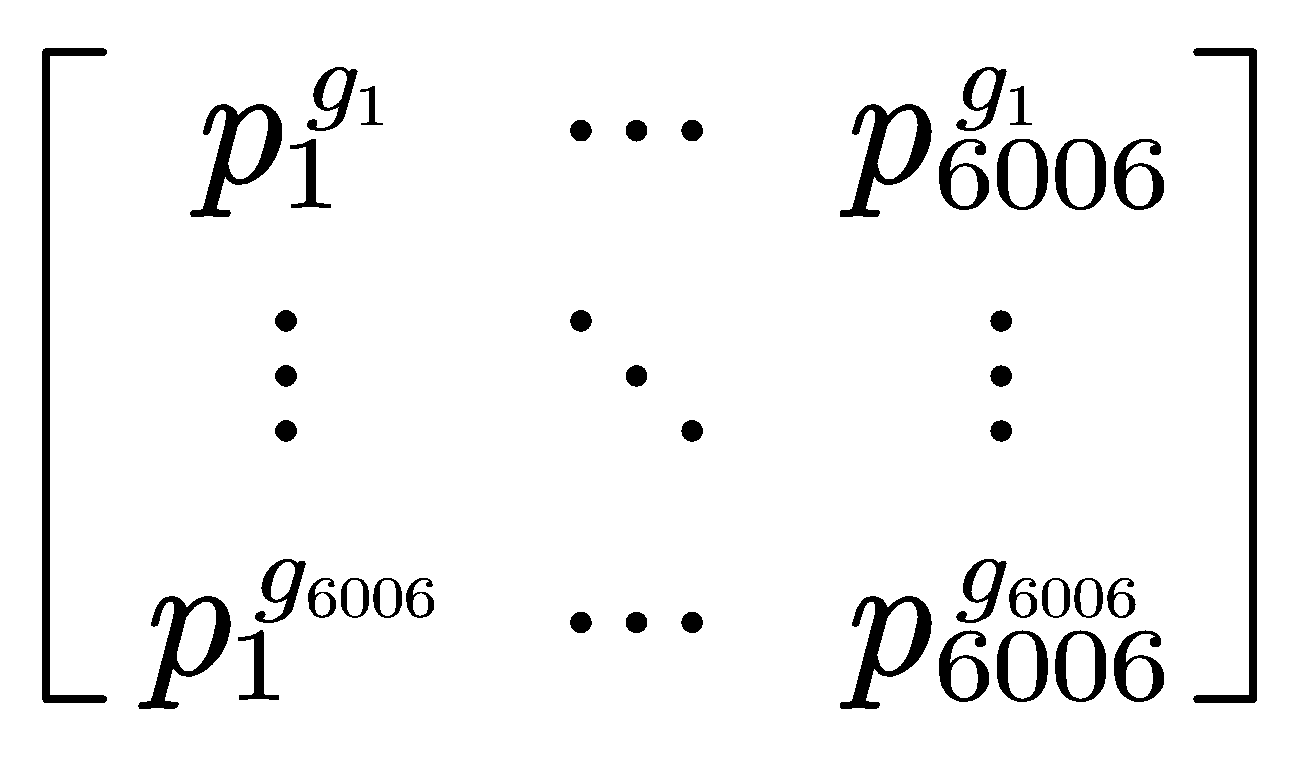
 and
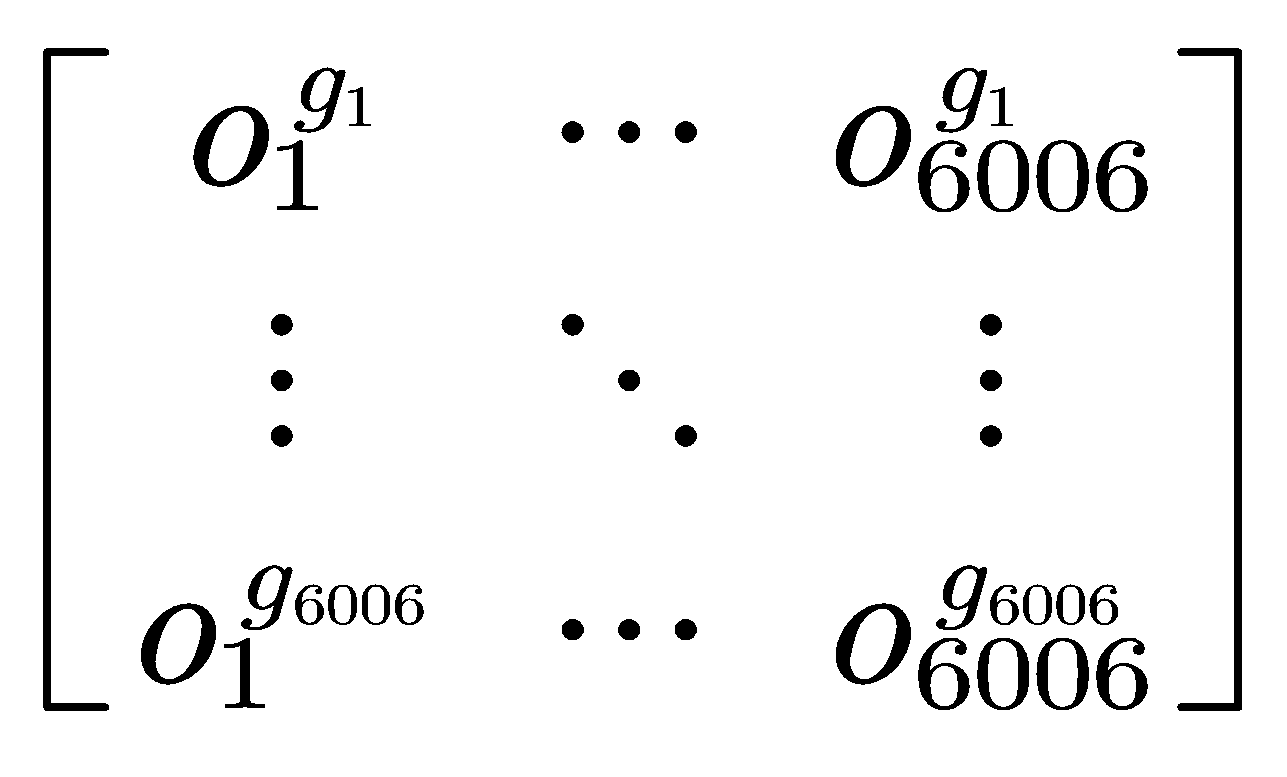

Where

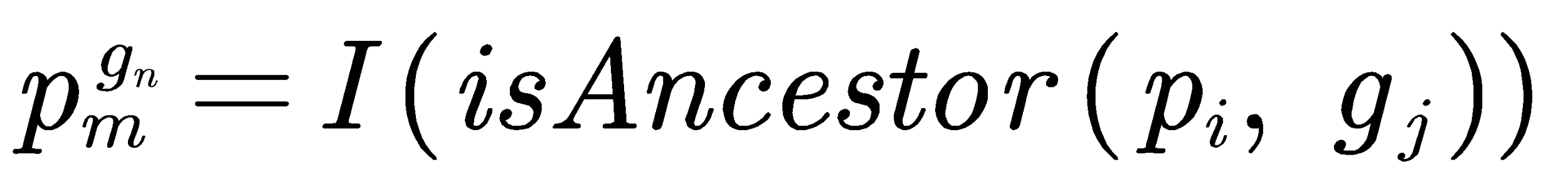

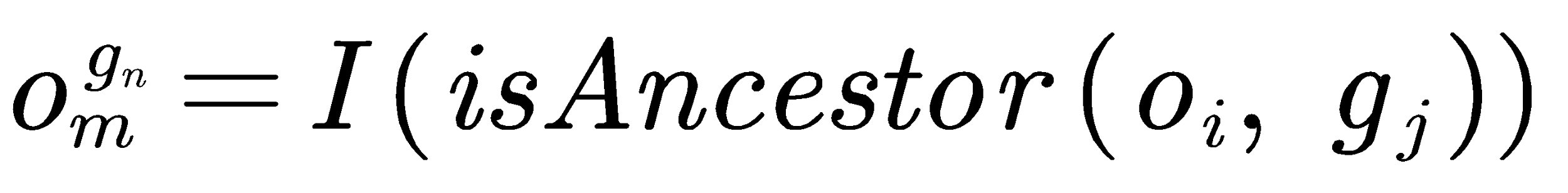

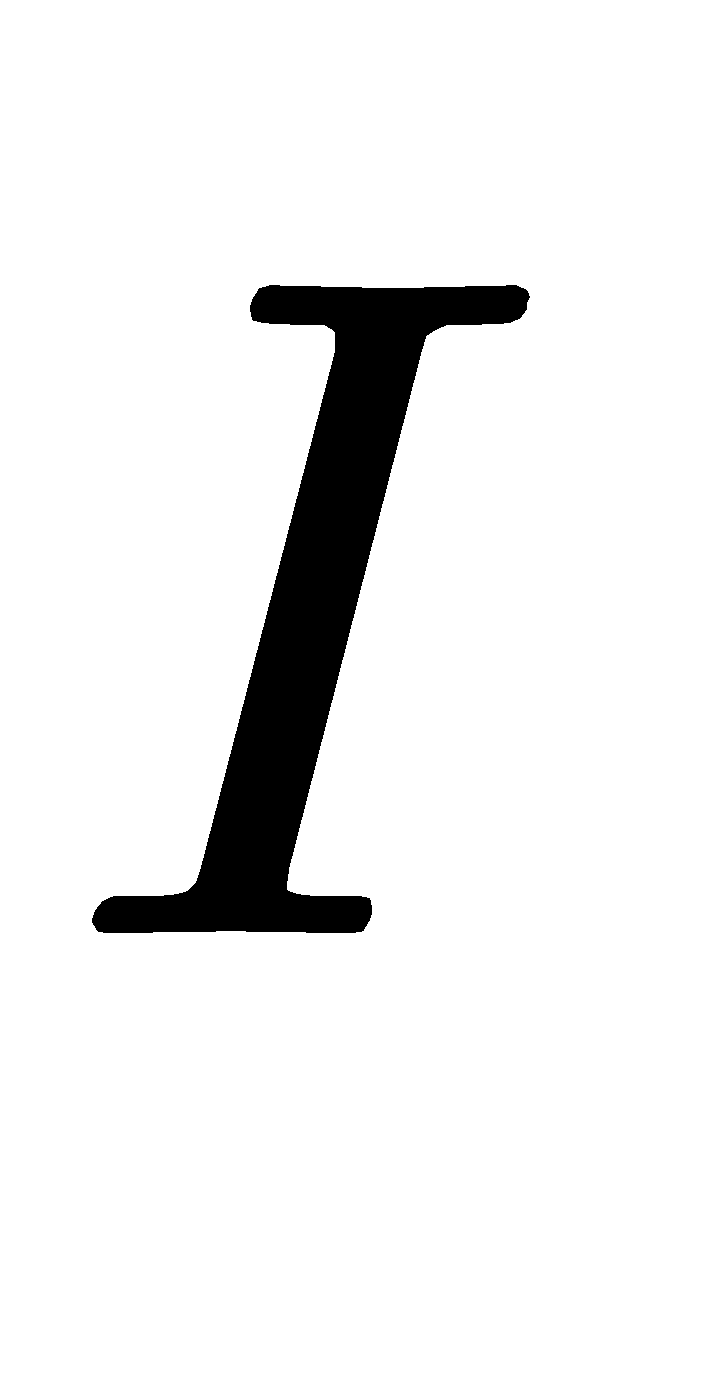
 is 1 when
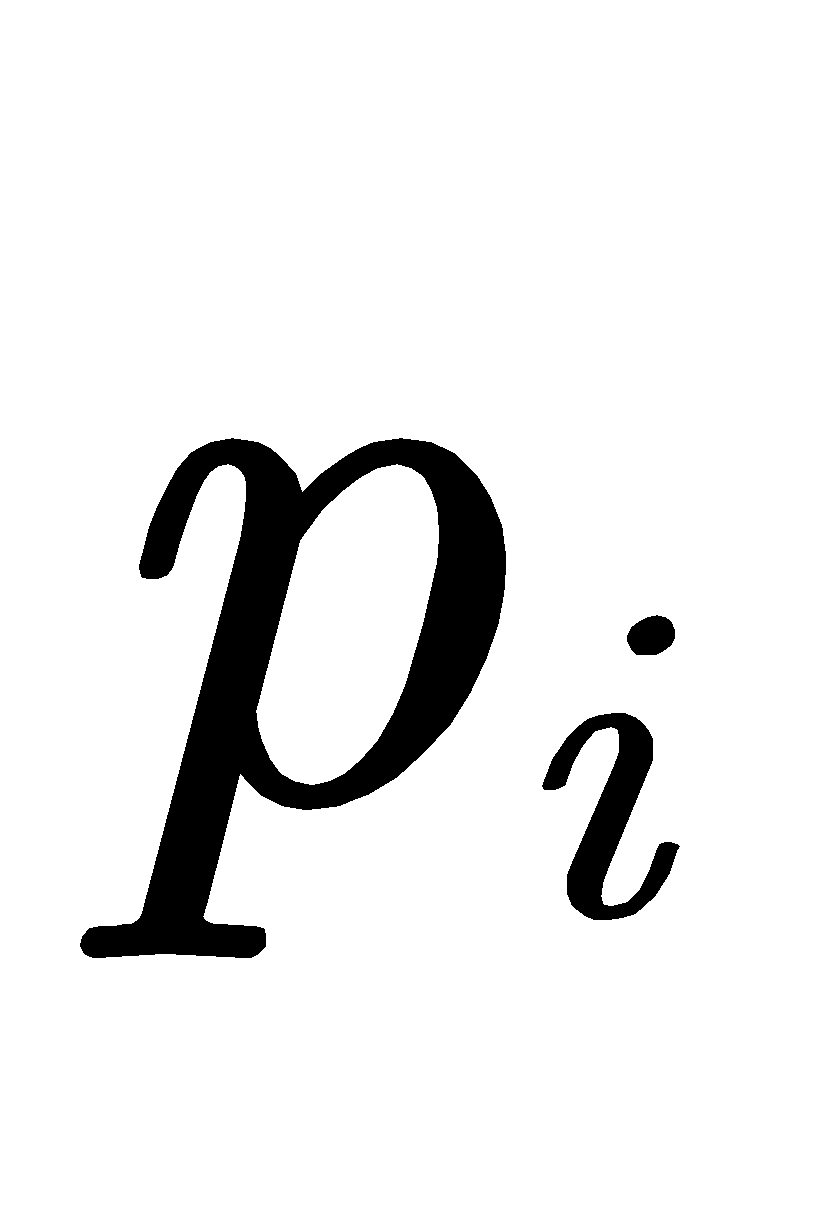
/
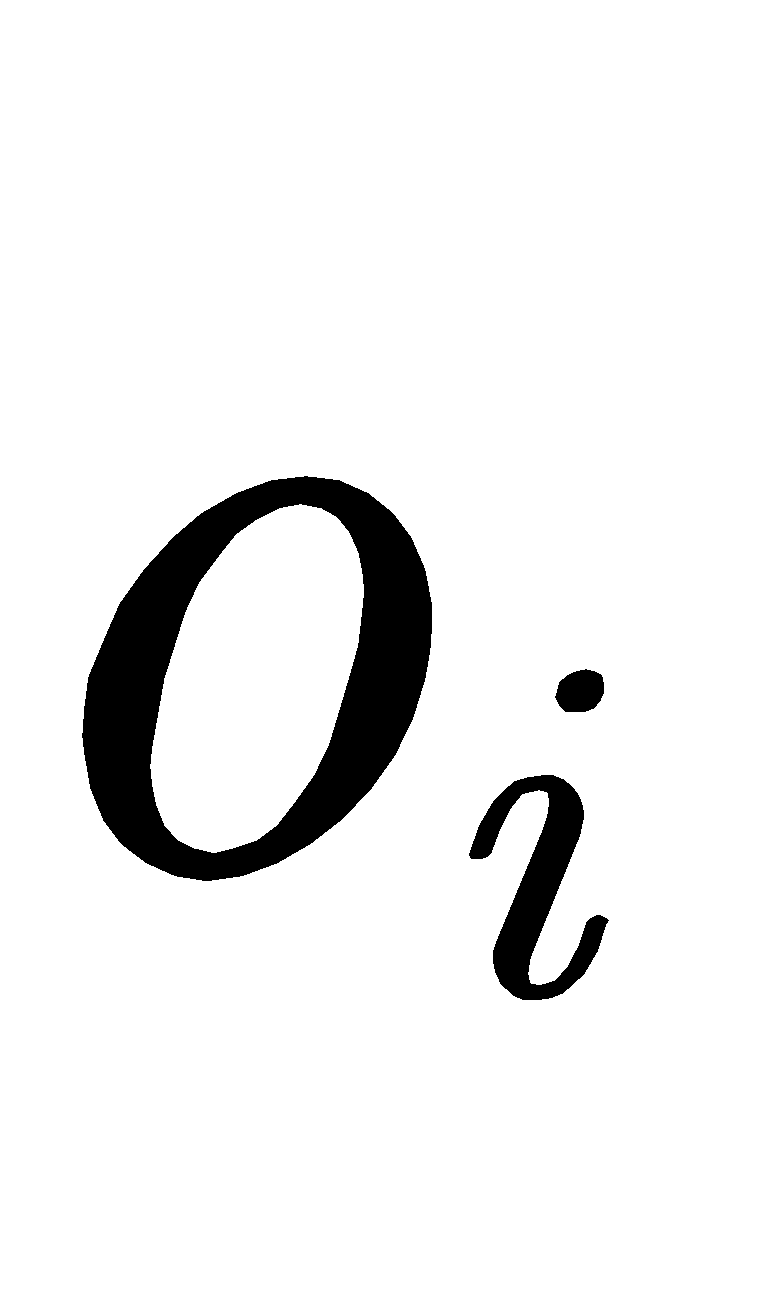
 is an ancestor of
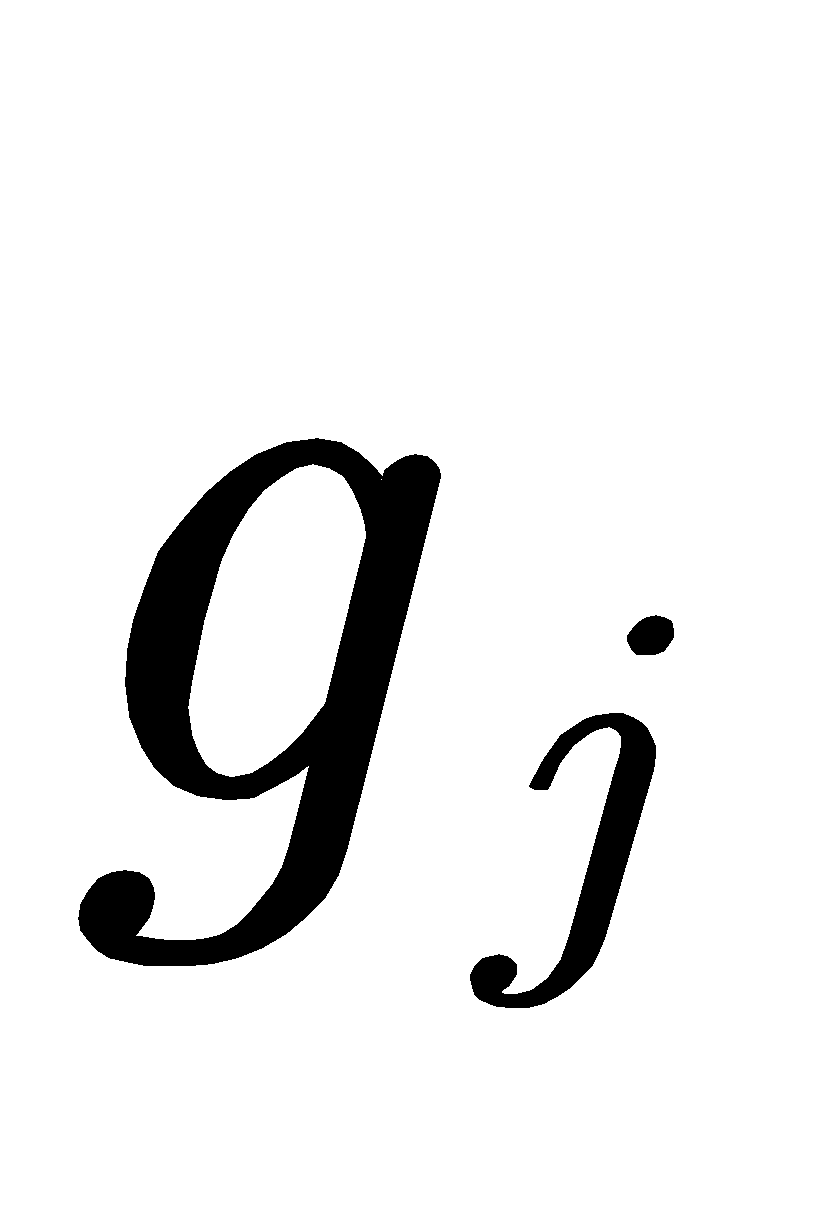
, otherwise 0. Within the matrix, each row represents a genus, each column represents a phylum or order, and each element represents whether the abundance of a genus should be aggregated into a phylum or order. Then, the total abundance for a dataset at phylum or order level could be calculated using simple matrix multiplication. For instance, the abundance at phylum level could be obtained through:

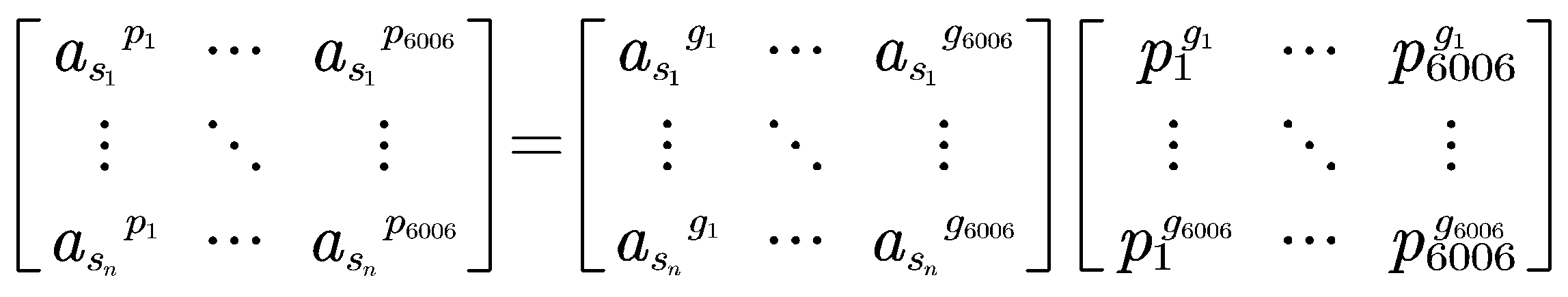

Such a process is implemented on the top of TensorFlow^4^ and thus could be automatically accelerated using multiple CPU cores or even a more advanced GPU.

### Supplementary Note 5

**Supplementary Note 5. Detailed procedure of performance comparison**

#### *Comparison with FEAST*

For EXPERT, we considered all biomes as sources, but only considered the seven independent biomes for comparison. We repeated a random cross-validation process five times. For each time of the cross-validation, we randomly selected 10% of the dataset as queries, and evaluated potential microbial sources by considering the remaining 90% of the dataset as sources. We enabled “fine-tuning” and “update statistics” options when optimizing the model. For other hyper-parameters used for training, please refer to **Supplementary Note 2** for a detailed description.

For FEAST, we considered seven independent biomes (**Supplementary Fig. S4**) as sources. After selecting 700 samples of these biomes as queries, we randomly selected 70, 140, and 120 community samples from the remaining samples (10, 20, and 30 samples for each biome).

#### *Comparison with pure NN approach*

We considered different proportions (1-10% by step size of 1%, and 10-90% by step size of 10%) of the human dataset as source samples, and quantified their contributions to pre-separated 10% of the dataset. For each setting of the proportion, the random selection of source samples and estimation was repeated five times. As a result, there were 90 (18 × 5) experiments performed. The source environment hierarchy is constructed by referring to the hierarchical biome classification from MGnify database^3^ and the ecosystem classification paths from GOLD database^32^.

### Supplementary Note 6

**Supplementary Note 6. Context-aware microbial source tracking**

#### *Infant succession study*

To demonstrate EXPERT’s utility in estimating the microbiota restoration, we considered the infant data from Backhed et al. First, we considered the samples from mothers, infants at birth, 4 months of age as sources, used them to construct a Transfer (HM) model, and estimated the source proportions for the samples at 12 months of age.

We repeated a random cross-validation process five times. For each time of the cross-validation, after grouping these samples by their delivery modes and sampling time, we randomly selected 10% of each group as queries, and evaluated potential microbial sources by considering the remaining 90% of each group as sources. We enabled “fine-tuning” and “update statistics” options when optimizing the model.

#### *Multi-disease study*

To characterize the disease patterns of human gut microbial communities, we repeated a random cross-validation process. For each time of the cross-validation, we used the human model as a fundamental model, and considered 90% of the dataset as sources and the remaining samples as queries. We enabled “fine-tuning” and “update statistics” options when optimizing the Transfer (HM) model. We also used the performance of the disease model to validate the disease pattern.

#### *Colorectal cancer study*

To characterize the progression of colorectal cancer using human gut microbial communities, we repeated a random cross-validation process. For each time of the cross-validation, we used the disease model and the human model as fundamental models, and considered 90% of the dataset as sources and the remaining samples as queries. We enabled “fine-tuning” and “update statistics” options when optimizing the Transfer (DM) and Transfer (HM) models. We also implemented an Independent model solely using the data in this context, to compare its performance with the two transferred models.
